# *In vivo* expansion of gene-targeted hepatocytes through transient inhibition of an essential gene

**DOI:** 10.1101/2023.07.26.550728

**Authors:** Marco De Giorgi, So Hyun Park, Adam Castoreno, Mingming Cao, Ayrea Hurley, Lavanya Saxena, Marcel A. Chuecos, Christopher J. Walkey, Alexandria M. Doerfler, Mia N. Furgurson, M. Cecilia Ljungberg, Kalyani R. Patel, Sarah Hyde, Tyler Chickering, Stephanie Lefebvre, Kelly Wassarman, Patrick Miller, June Qin, Mark K. Schlegel, Ivan Zlatev, Rich Gang Li, Jong Kim, James F. Martin, Karl-Dimiter Bissig, Vasant Jadhav, Gang Bao, William R. Lagor

**Author notes:** Corresponding authors Correspondence: William R. Lagor, Ph.D., Department of Integrative Physiology, Baylor College of Medicine, Houston, TX 77030, USA. Gang Bao, Ph.D., Department of Bioengineering, Rice University, Houston, TX 77030, USA.

## Abstract

Homology Directed Repair (HDR)-based genome editing is an approach that could permanently correct a broad range of genetic diseases. However, its utility is limited by inefficient and imprecise DNA repair mechanisms in terminally differentiated tissues. Here, we tested “Repair Drive”, a novel method for improving targeted gene insertion in the liver by selectively expanding correctly repaired hepatocytes *in vivo*. Our system consists of transient conditioning of the liver by knocking down an essential gene, and delivery of an untargetable version of the essential gene *in cis* with a therapeutic transgene. We show that Repair Drive dramatically increases the percentage of correctly targeted hepatocytes, up to 25%. This resulted in a five-fold increased expression of a therapeutic transgene. Repair Drive was well-tolerated and did not induce toxicity or tumorigenesis in long term follow up. This approach will broaden the range of liver diseases that can be treated with somatic genome editing.

## Introduction

The liver is the metabolic hub of the body, and is readily amenable to gene delivery with viral and non-viral vectors such as adeno-associated viruses (AAV) and lipid nanoparticles (LNP). Somatic genome editing trials in humans with CRISPR/Cas9 and Base Editors are already underway with promising results.^1, 2^ However, these are aimed at disruption of genes (i.e. *TTR*, *PCSK9*) which is feasible for only a very small subset of genetic targets. The vast majority of liver diseases require *correction* rather than disruption. Base editors^3, 4^ and prime editors^5^ can install precise changes. However, not all sites are targetable, and customization of reagents for each private mutation is a daunting logistical hurdle for clinical application. Homology directed repair (HDR) can make very precise changes, including large and small insertions, deletions, and replacements of entire genes. However, its utility is severely limited by the requirement for active cell division.^6, 7^

Numerous strategies are being pursued to improve the efficiency of HDR. Small molecule drugs can promote HDR or inhibit non-homologous end joining (NHEJ) repair – the predominant and error-prone DNA repair mechanism in non-dividing cells.^8–10^ However, global inhibition of DNA repair pathways carries the risk of cytotoxicity and additional mutations. More promising approaches leverage the inherent regenerative capacity of the liver. Nygaard et al. targeted transgenes to the Albumin (*Alb)* locus carrying a short hairpin RNA (shRNA) against 4-OH phenylpyruvate dioxygenase (*Hpd*), whose product catalyzes an early step in tyrosine catabolism. This conferred a growth advantage to gene-modified hepatocytes upon inhibition of fumarylacetoacetate hydrolase (Fah), allowing cells expressing the therapeutic transgene to survive and repopulate the liver.^11^ Similar methods using microRNA or gene editing to alter the diphtheria toxin receptor have been used to select for human cells in murine xenograft models.^12, 13^ More recently, Vonoda et al. used CRISPR/Cas9 to permanently inactivate NADPH-cytochrome p450 reductase (*Cypor*) - an essential cofactor for cytochrome P450 (*Cyp*).^14^ *Cypor* deletion gave gene-modified hepatocytes resistance to toxic concentrations of the Cyp substrate acetaminophen, forcing their expansion upon exposure to the drug.^14, 15^ In each case, expression of therapeutic transgenes can be coupled to a genetically encoded survival advantage. These studies demonstrate that the plasticity of the liver can be leveraged in unique ways to increase the proportion of HDR-targeted cells. However, all reported methods to date involve the use of toxic compounds, or permanently alter liver metabolism. Translation to human gene therapy will require methods that can be transiently applied, and restore normal liver physiology following selection.

Here, we report Repair Drive, an approach to selectively expand gene*-*targeted hepatocytes *in vivo*. Repair Drive involves the transient inhibition of an essential gene with small interfering RNA (siRNA) to gradually deplete incorrectly targeted cells. HDR-repaired hepatocytes express an untargetable version of the essential gene *in cis* with the therapeutic transgene. Over time, correctly repaired cells divide and repopulate the liver. Expression of the essential gene is restored to normal following the selection period. Most importantly, the proportion of correctly repaired hepatocytes is dramatically increased, enabling durable therapeutic transgene expression.

## Results

### Selective expansion of targeted hepatocytes through inhibition of an essential gene

To increase the efficiency of HDR, we developed a strategy to selectively expand gene-targeted hepatocytes *in vivo* – referred to as Repair Drive (Fig. 1). Repair Drive consists of (1) transient conditioning of the liver by knocking down an essential gene using N-Acetylgalactosamine (GalNAc)-conjugated siRNA^16^ (Fig. 1a); and (2) delivery of an untargetable version of the essential gene (as a selectable marker) *in cis* with the therapeutic transgene to a highly expressed liver locus (Apolipoprotein A1 - *Apoa1*) (Fig. 1b).^17^ We previously showed that the *Apoa1* locus can drive strong expression of therapeutic transgenes in the liver.^17^ This should result in gradual depletion of untargeted hepatocytes during liver conditioning and a selective expansion of targeted cells, which are protected by the selectable marker and express the therapeutic transgene (Fig. 1c). As conditioning agent, we exploited the gene *Fah*, which is essential for hepatocytes. Fah catalyzes the last step in tyrosine catabolism, and its loss results in accumulation of toxic catabolites, leading to cell death (Extended Data Fig. 1a).^18^ Following an *in vitro* screening of siRNAs targeting murine *Fah*, we tested the top two for potency *in vivo* (Extended Data Fig. 1). *Fah*-siRNA6 produced more efficient and durable knockdown than siRNA10 (Extended Data Fig. 1d) and was therefore selected for Repair Drive (hereinafter referred as *Fah*-siRNA).

**Fig. 1.**
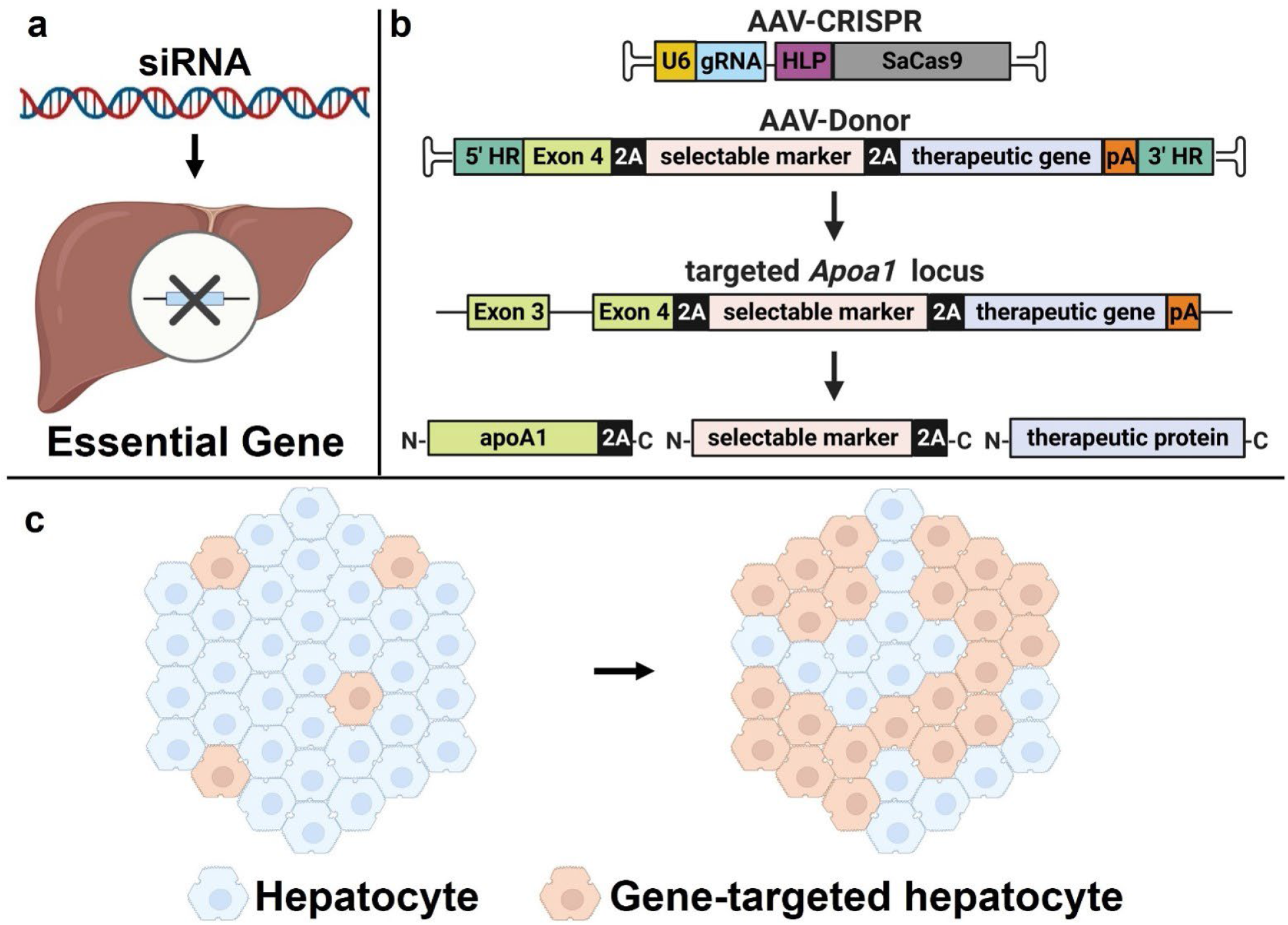
Repair Drive protocol. **a**, Liver conditioning: the liver is treated with a GalNAc-conjugated siRNA for transiently knocking down an essential gene. **b**, Targeting strategy: a dual AAV system is used for transgene targeted integration into the *Apoa1* locus. AAV-CRISPR encodes a gRNA targeting the 3’ untranslated region (3’UTR) of *Apoa1* - under the control of U6 promoter - and the *Staphylococcus aureus* Cas9 (SaCas9) driven by a liver-specific promoter (HLP). AAV-Donor contains the final coding exon of *Apoa1* (exon 4) followed by the promoterless coding sequences of human *FAH* (as a selectable marker) and therapeutic transgene, all linked in frame by 2A ribosomal skipping sequences. pA: poly-adenylation signal; HR: homology region to the *Apoa1* locus. Following HDR integration of the donor cassette into the *Apoa1* locus, targeted hepatocytes express 3 separate proteins: apoA1 and the selectable marker - both with a C-terminal tail of ∼20 amino acids derived by the translation of 2A - and the therapeutic protein. **c**, Scheme of expansion: the liver conditioning agent transiently induces cell death of untargeted hepatocytes, which causes compensatory cell proliferation. In parallel, correctly targeted hepatocytes are resistant to liver conditioning through the selectable marker and expand over time, therefore boosting the expression of therapeutic transgenes. Created with BioRender.com.

To test Repair Drive, we injected adult mice with a dual AAV-CRISPR/Donor system to insert our selectable marker (human *FAH*, which is resistant to the *Fah*-siRNA sequence) in tandem with a fluorescent reporter (TdTomato) into the *Apoa1* locus (Fig. 1b and 2a). Following genome editing, mice were injected monthly with the *Fah*-siRNA for selection (Fig. 2a). Liver conditioning resulted in efficient knockdown of Fah (Extended Data Fig. 2), which caused transient hepatocyte death (Supplementary Fig. 1-3) and modest elevations of plasma alanine transaminase (ALT) (Fig. 2b and Supplementary Fig. 1b). In the CRISPR/Donor + siRNA group (hereinafter referred as Repair Drive mice), we observed enrichment of HDR-modified *Apoa1* alleles-up to 3±1.4% (Fig. 2c-d). In parallel, we observed a significantly decreased indel formation-30.2±2.6 vs. 37.3±5.6% in CRISPR/Donor only injected mice (hereinafter referred as Unselected mice) (Extended Data Fig. 2a), but no significant changes in NHEJ-mediated insertion of AAV genomes (Fig. 2e). Along with more HDR-modified *Apoa1* alleles, Repair Drive mice had increased expression of the 2A-tagged Apoa1 and FAH products (Extended Data Fig. 2 and Supplementary Fig. 1). Most importantly, Repair Drive mice showed a dramatic increase in TdTomato positive hepatocytes-9±3 vs. 0.3±0.3% in Unselected mice (Fig. 2f, g and Supplementary Fig. 1). These hepatocytes were arranged in colonies, suggesting the clonal expansion of corrected cells following siRNA treatment (Supplementary Fig. 4). In addition, a single-nucleus RNA sequencing (snRNA-seq) analysis revealed that the corrected cells clustered mostly in a hepatocyte subpopulation enriched for diploidy and portal zonation markers (Supplementary Fig. 5 and Tables S1-4).^19, 20^

**Fig. 2.**
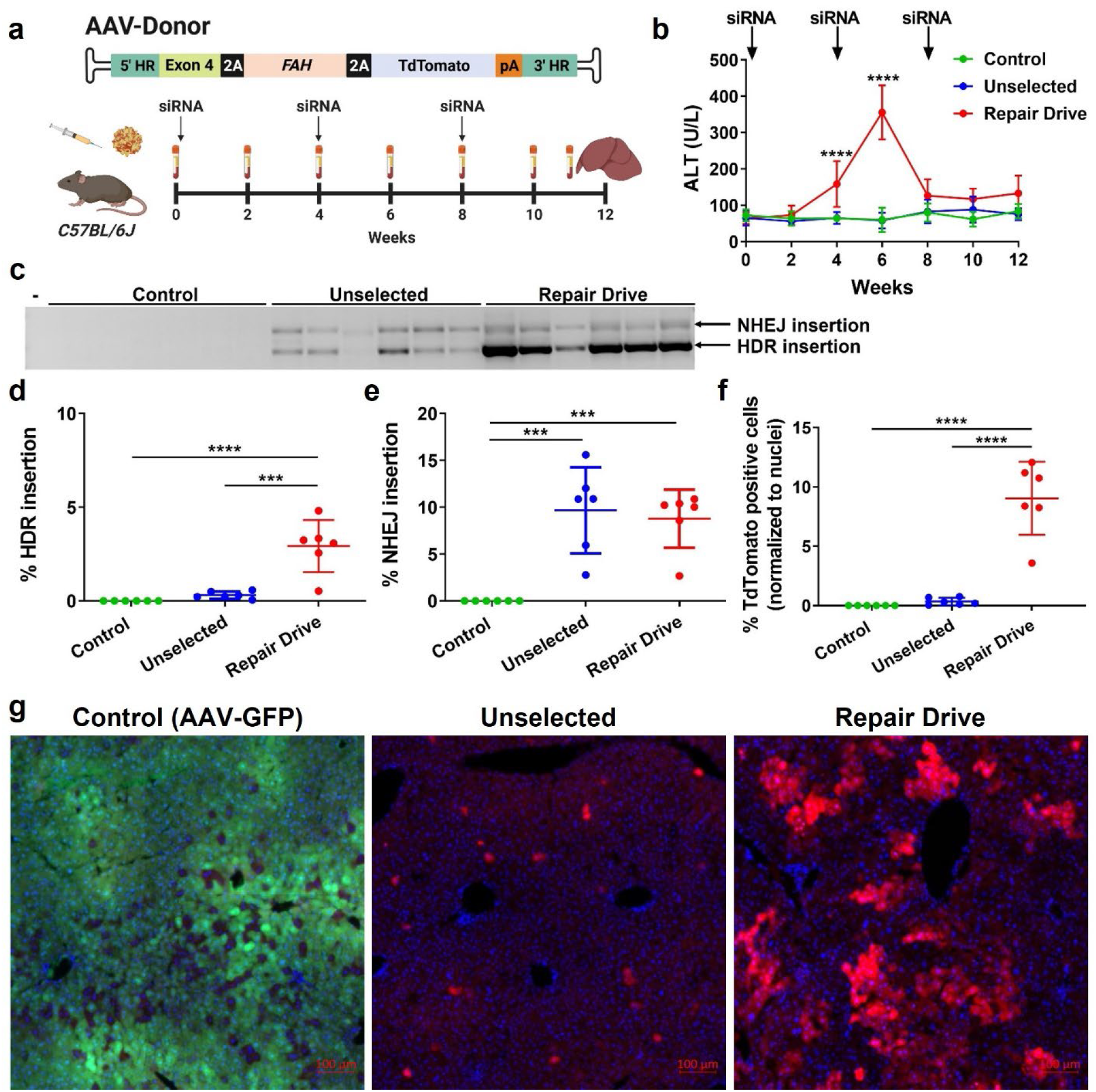
Expansion of *Apoa1*-targeted hepatocytes. **a**, An AAV-Donor was designed to integrate the promoterless coding sequences of human *FAH* (as a selectable marker) and TdTomato (as a reporter gene) into the *Apoa1* locus. Eight-week-old *C57BL/6J* mice were injected with 5×10^11^ GC of AAV-CRISPR and 1×10^11^ GC of AAV-Donor. Control mice were injected with 6×10^11^ GC of AAV-GFP. Following genome editing, mice were injected with 3 mg/kg of *Fah*-siRNA or saline every four weeks and sacrificed at twelve weeks post-AAV injection for liver analysis. Blood was collected before injection (time 0) and every two weeks for plasma analysis. Unselected: CRISPR/Donor only injected mice. Repair Drive: CRISPR/Donor + siRNA injected mice. Created with BioRender.com. **b**, Plasma ALT measurement showing transient elevation in Repair Drive mice. **c**, PCR from liver show integration of AAV-Donor into the *Apoa1* locus. Two main products were observed: correct HDR (886 bp) and NHEJ insertion (1778 bp). The forward primer binds to the *Apoa1* locus upstream of the 5’ HR and reverse primer binds to the *FAH* coding sequence. Minus (-) indicates water-only control. **d**, ddPCR showing the frequency of *Apoa1* allele with HDR-mediated integration of AAV-Donor. **e**, ddPCR showing the frequency of *Apoa1* allele with NHEJ insertion of ITR sequences. **f**, Quantification of TdTomato-positive hepatocytes relative to total nuclei per field. **g**, Representative direct fluorescence microscopy showing GFP and TdTomato respectively in Control and AAV-CRISPR/Donor-injected mice subjected or no to Repair Drive. Scale bar is 100 µm. Data are expressed as mean ± standard deviation (s.d.) (n= 6 mice per group), with significance determined by one-way and two-way analysis of variance (ANOVA) followed by Tukey test, respectively in **d**-**f** and **b**. *** p<0.001 and **** p<0.0001. In (**b**), **** p<0.0001 Repair Drive vs. Control and Unselected mice.

**Fig. 3.**
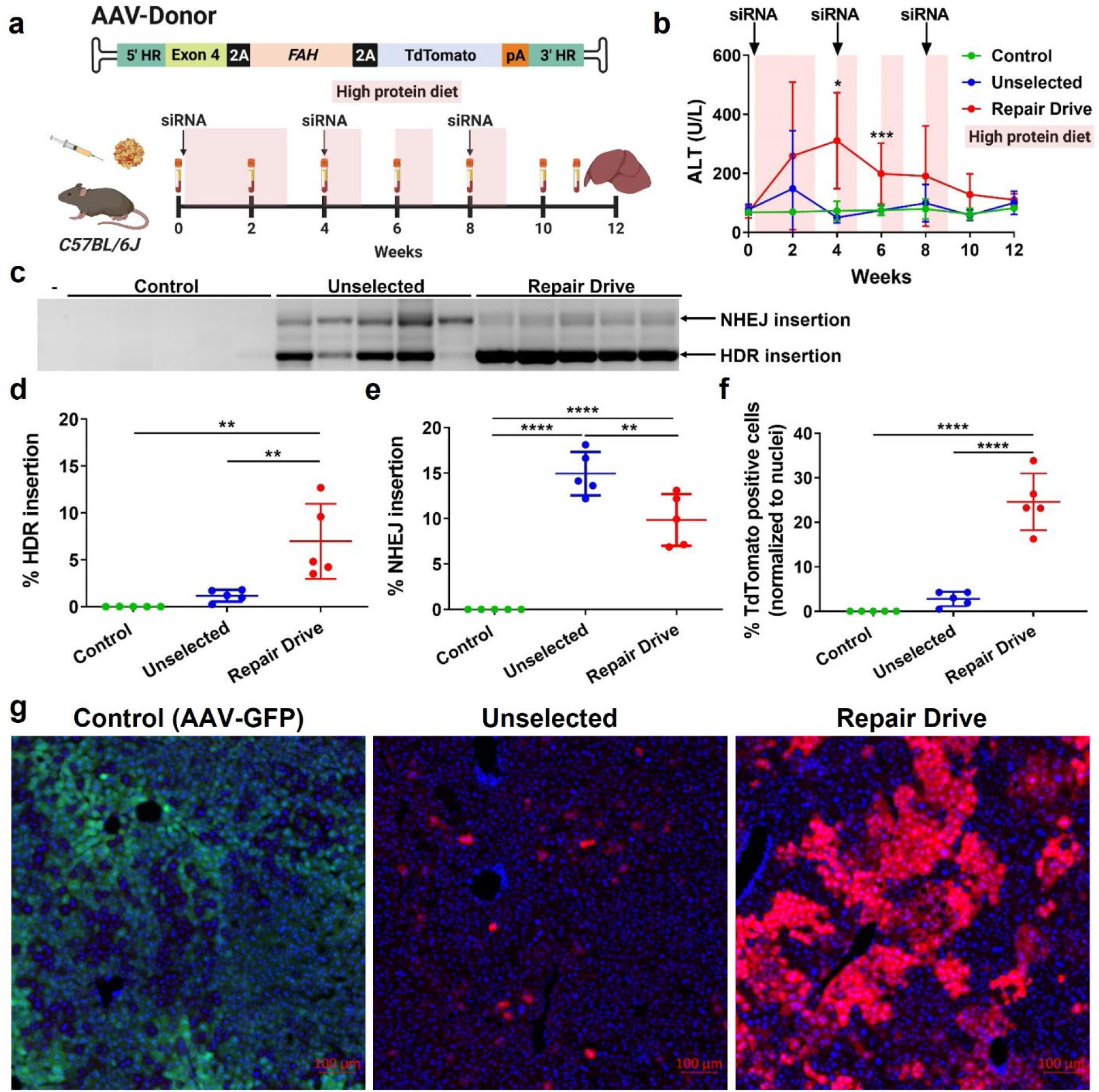
Increased expansion of *Apoa1*-targeted hepatocytes with high protein diet. **a**, Eight-week-old *C57BL/6J* mice were injected with AAVs (5×10^11^ GC of AAV-CRISPR and 1×10^11^ GC of AAV-Donor or 6×10^11^ GC of AAV-GFP as control) and subjected to the expansion protocol with monthly injection of *Fah*-siRNA (3 mg/kg or saline as control). In addition, mice were cycled on and off with a high protein diet (on-periods shown in pink). Blood was collected at time 0 and every two weeks and mice were sacrificed at twelve weeks post AAV-injection. Created with BioRender.com. **b**, Plasma ALT measurement showing transient elevation in Repair Drive mice. **c**, PCR from liver show integration of AAV-Donor into the *Apoa1* locus. Two main products were observed: correct HDR (886 bp) and NHEJ insertion (1778 bp). The forward primer binds to the *Apoa1* locus upstream of the 5’ HR and reverse primer binds to the *FAH* coding sequence. Minus (-) indicates water-only control. **d**, ddPCR showing the frequency of *Apoa1* allele with HDR-mediated integration of AAV-Donor. **e**, ddPCR showing the frequency of *Apoa1* allele with NHEJ insertion of ITR sequences. **f**, Quantification of TdTomato-positive hepatocytes relative to total nuclei per field. **g**, Representative direct fluorescence microscopy showing GFP and TdTomato respectively in Control and AAV-CRISPR/Donor-injected mice subjected or no to Repair Drive. Scale bar is 100 µm. Data are expressed as mean ± s.d. (n= 5 mice per group), with significance determined by one-way and two-way ANOVA followed by Tukey test, respectively in **d**-**f** and **b**. * p<0.05, ** p<0.01, *** p<0.001 and **** p<0.0001. In (**b**), * p<0.05 Repair Drive vs. Control mice and *** p<0.001 Repair Drive vs. Control and Unselected mice.

### Increased selective expansion with high-protein diet

Next, we sought to increase the selective pressure of the liver conditioning. To do this, mice were injected with AAVs and *Fah*-siRNA as described above and then cycled with a high-protein diet to increase tyrosine catabolism and hence the accumulation of toxic metabolites in the liver (Fig. 3a). This conditioning protocol resulted in transient ALT elevation and body weight loss, which fully resolved by the end of the experiment (Fig. 3b and Supplementary Fig. 6). At the endpoint, we observed robust knockdown of murine Fah (Extended Data Fig. 3) and evidence of cell death (Supplementary Fig. 7) in Repair Drive livers. We also detected around 20- and 14-fold lower AAV-CRISPR and AAV-Donor genome copies (GC) in Repair Drive livers as compared to Unselected (Supplementary Fig. 8). Most importantly, we observed a 6-fold increase in HDR-corrected *Apoa1* allele-7±4 vs. 1.2±0.6% in Unselected mice-(Fig. 3c, d) and significant decreases of both indel formation-32±5 vs. 38.8±4.1%, and NHEJ-mediated insertion of AAV genomes– 9.8±2.8 vs. 14.9±2.4% (Extended Data Fig. 3a and Fig. 3e). The enrichment of HDR-corrected *Apoa1* alleles resulted in strong expression of the 2A-tagged Apoa1 and FAH products, which did not affect endogenous Apoa1 levels (Extended Data Fig. 3). The rate of TdTomato positive hepatocytes dramatically increased 8-fold in Repair Drive mice as compared to Unselected mice, reaching 24.6±6.4 vs. 2.8±1.6% (Fig. 3f, g). The positive hepatocytes were arranged in colonies, which were larger than what was observed in the previous experiment (Supplementary Fig. 9).

### Characterization of on-target editing events

To accurately quantify comprehensive gene editing outcomes introduced at the *Apoa1* locus, we combined long-range PCR with single-molecule real-time sequencing (SMRT-seq) (Supplementary Fig. 10-15 and Table S5).^21^ Long-read sequencing showed the presence of *Apoa1* alleles carrying heterogeneous integration events of AAV-Donor (Fig. 4) and AAV-CRISPR (Supplementary Fig. 14-15). Repair Drive livers showed a significant decrease of small indels and NHEJ-insertion of AAV genomes as compared to Unselected (Fig. 4a-d and Supplementary Fig. 13). About 80% of these NHEJ-insertions contained inverted terminal repeats (ITRs) and mapped to AAV-CRISPR (Supplementary Fig. 13-14). We also found full AAV genome capture or AAV concatemers with both head-to-tail or tail-to-tail configurations of AAV-CRISPR and AAV-Donor (Supplementary Fig. 15). Most importantly, a dramatic increase in HDR integration of AAV-Donor was observed in Repair Drive mice as compared to Unselected (Fig. 4a-d and Supplementary Fig. 13). The HDR reads were categorized into four subtypes: (1) HDR at both the 5’ and 3’ ends (HDR), (2) 5’HDR with 3’NHEJ (5’HDR-3’NHEJ), (3) HDR with partial truncation of TdTomato - HDR(-Td) - and (4) 5’HDR(-Td)-3’NHEJ (Fig. 4e). The frequency of HDR and 5’HDR(-Td) events increased to a greater extent than the other subtypes following expansion (Fig. 4b and d). All the HDR insertions were expected to result in functional expression of the selectable marker *FAH* (Fig. 4e).

**Fig. 4.**
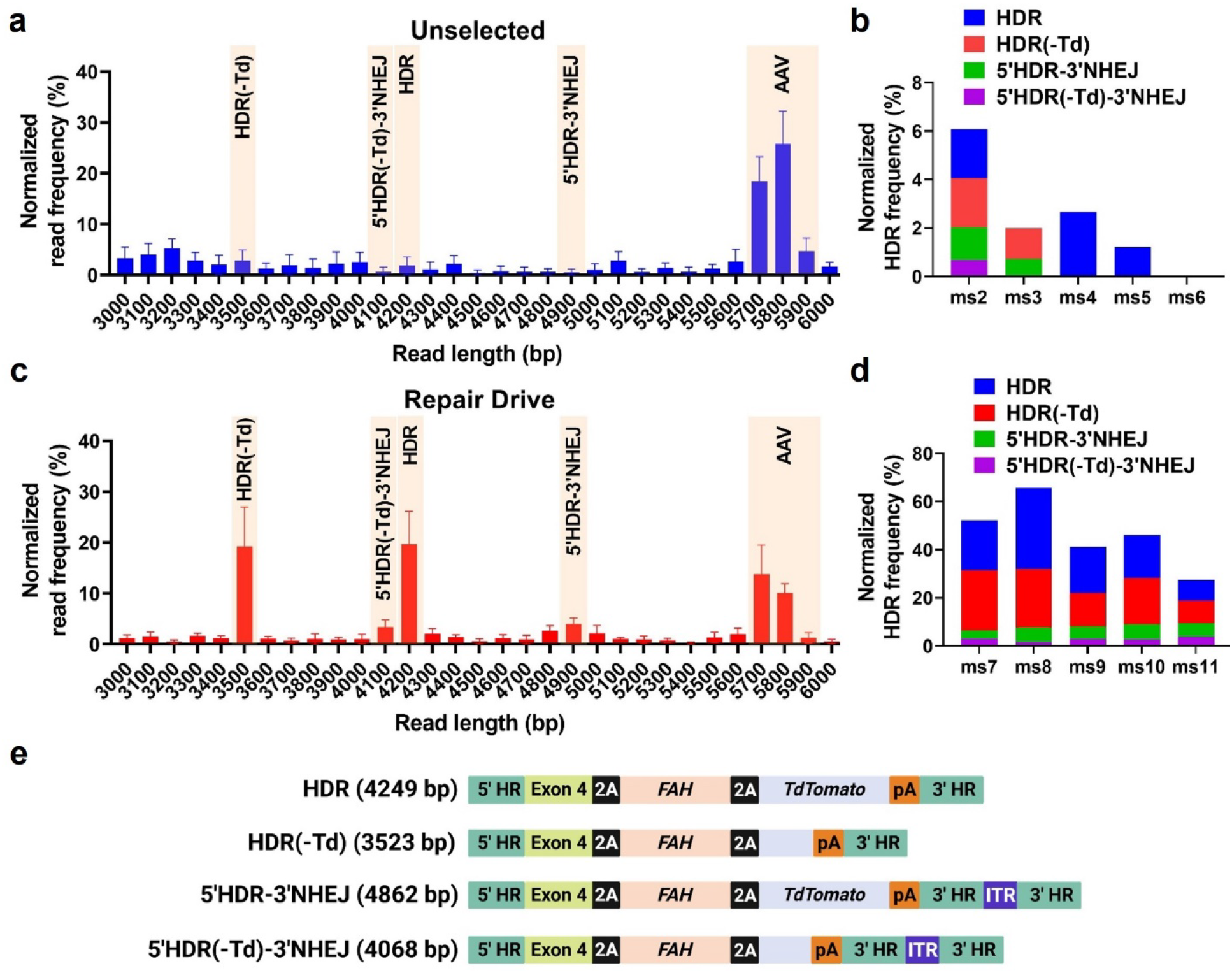
Selective enrichment of HDR integration events at the *Apoa1* locus with Repair Drive. **a**, Normalized frequency distribution of sequences with 3000 - 6000 bp read length in Unselected mice. Highlighted the reads corresponding to different HDR subtypes - HDR(-Td) (∼3500 bp), 5’HDR(-Td)-3’NHEJ (∼4100 bp), HDR (∼4200 bp) and 5’HDR-3’NHEJ (∼4900 bp) and the reads corresponding to NHEJ-mediated AAV genome insertions (∼5700 - 5900 bp). **b**, Normalized frequency of HDR subtypes in Unselected mice. Ms: mouse. **c**, Normalized frequency distribution of sequences with 3000 - 6000 bp read length in Repair Drive mice. Highlighted the reads corresponding to different HDR subtypes - HDR(-Td) (∼3500 bp), 5’HDR(-Td)-3’NHEJ (∼4100 bp), HDR (∼4200 bp) and 5’HDR-3’NHEJ (∼4900 bp) and the reads corresponding to NHEJ-mediated AAV genome insertions (∼5700 - 5900 bp). **d**, Normalized frequency of HDR subtypes in Repair Drive mice. Ms: mouse. **e**, Schematics of the four subtypes of HDR integration outcomes expected to result in *Apoa1*-driven transgene expression. **e** Created with BioRender.com.

### Off-target integration of AAV-Donor

The integration of AAV-Donor in genomic sites other than *Apoa1* (off-target) could confer a growth advantage independent of downstream transgene expression. To address this, we developed a <u>G</u>enome-wide Integration <u>S</u>ite of <u>A</u>AV-Donor identified by <u>seq</u>uencing assay (GISA-seq) for unbiased genome-wide retrieval of AAV-Donor integration sites, using primers amplifying outward from the TdTomato sequence (Fig. 5a, Supplementary Fig. 16-22 and Tables S6-8). Most reads mapped to episomal AAV-CRISPR and AAV-Donor (respectively 81.7±2.2 vs. 73.4±1.9%, and 15.6±3.1 vs. 10.3±1.3% in Unselected vs. Repair Drive livers) (Fig. 5b). The reads in AAV-CRISPR were a result of tail-to-tail and tail-to-head rearrangements with AAV-Donor (Supplementary Fig. 18). In Unselected livers, only 2.6±1% of reads mapped to the mouse genome (Fig. 5b). ∼28% of these mapped to the *Apoa1* locus, whereas the remainder annealed to off-target sites - ∼60% including ITR (Fig. 5c-e and Supplementary Fig. 18-20). The frequency of reads with genome junction increased of ∼6-fold in Repair Drive mice as compared to Unselected (16.3±3%, Fig. 5b), with ∼80% of them mapping to the *Apoa1* locus (Fig. 5c). The frequency of off-target integration dramatically decreased in Repair Drive mice relative to Unselected mice – 17.8±10.8 vs. 59.9±20.6% with ITR and 4.1±08 vs. 12.2±4.6% without ITR (Fig. 5c-e and Tables S7-8). The off-target sites were distributed across the whole genome but showed a slightly higher preferential distribution in chromosomes 4 and 5 in Repair Drive mice (Fig. 5f). AAV-Donor integrated mainly in intergenic and intronic regions, with a preferential integration into introns in Repair Drive mice (Fig. 5g). Interestingly, we observed redundant integration in previously reported AAV integration hotspots - *Gm10800*, *Alb*, Argininosuccinate synthase 1 (*Ass1*) and Alpha-fetoprotein (*Afp*) (Fig. 5h,i).^22, 23^ *Alb* showed different unique integration sites and one hotspot in intron 12 (Supplementary Fig. 21). In Repair Drive livers, we found fewer unique integration sites in *Alb* but higher counts for sites sharing the same integration junction, potentially derived by clonal expansion of *Alb*-targeted cells (Supplementary Fig. 21). In addition to *Alb* and *Ass1*, we observed off-target integration in other highly expressed loci in the liver, such as Carbamoyl-phosphate synthase 1 (*Cps1*), ATP binding cassette subfamily C member 2 (*Abcc2*) and Tetratricopeptide Repeat Domain 39C (*Ttc39c*) (Fig. 5h,i and Supplementary Fig. 21-22).

**Fig. 5.**
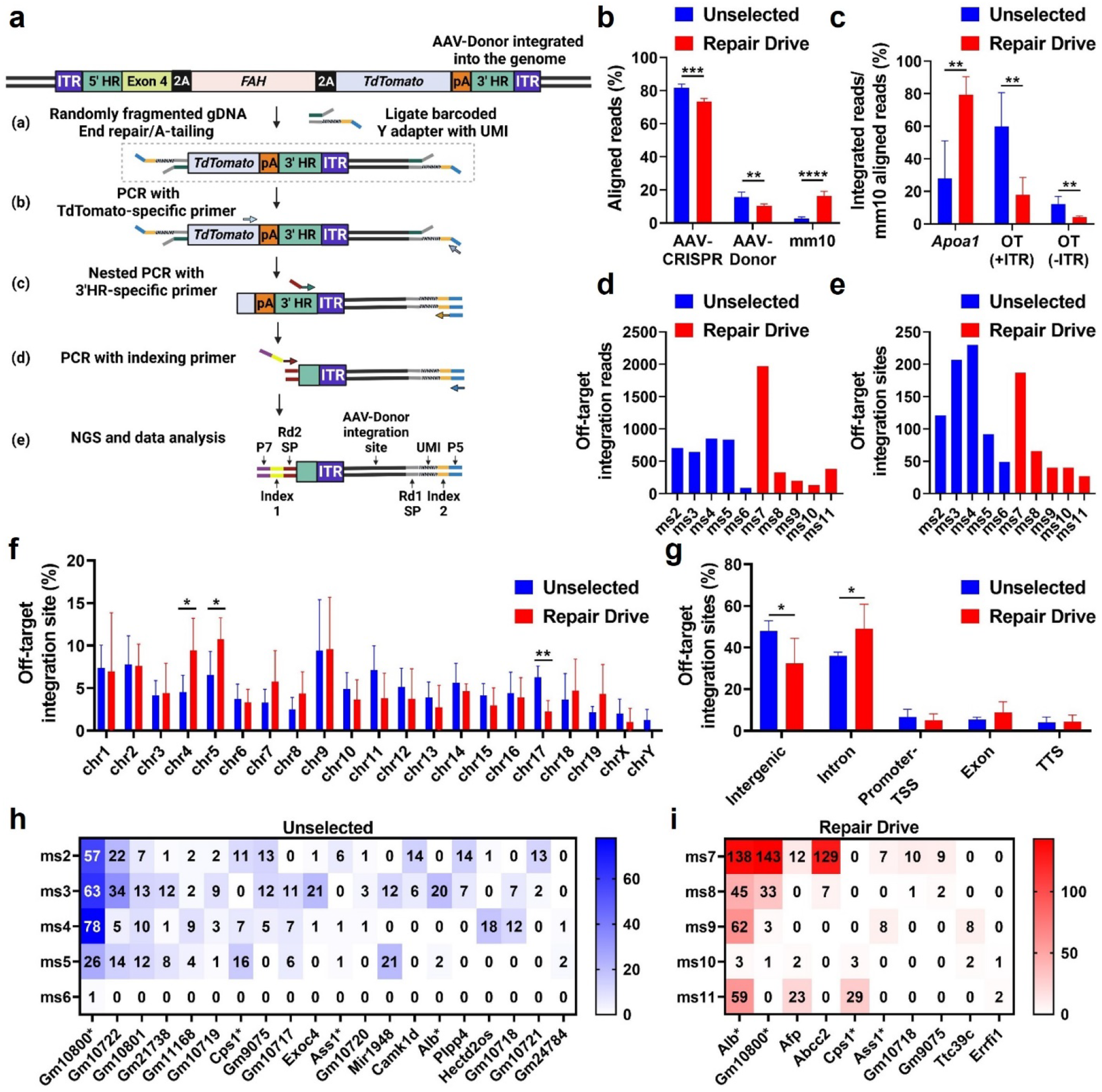
Analysis of genome wide AAV-Donor integration. **a**, Schematics of GISA-seq for genome-wide determination of AAV-Donor integration sites. The assay is based on anchored PCR reactions for amplifying outward from the AAV-Donor to sequence across junctions involving genome integration. (a) Genomic DNA was randomly fragmented and subsequently processed for end-repair, A-tailing, and ligation of unique Y-adapters containing Unique Molecular Identifier (UMI) and sample-specific P5 index. (b) Y-adapter-ligated DNA fragments were amplified by PCR using a TdTomato-specific forward primer and a reverse primer binding to the P5 sequence on Y-adapter. (c) Subsequent nested PCR was performed using a staggered forward primer annealing at 3’HR and sample-specific P5 index reverse primer. (d) Sample-specific P7 index was added by PCR. (e) Final PCR product for next-generation sequencing and data analysis. Rd1 SP: Read1 sequencing primer. Rd2 SP: Read2 sequencing primer. Created with BioRender.com. **b**, Frequency of reads aligned to episomal AAV-CRISPR and AAV-Donor, and mouse genome (mm10) relative to total reads. **c**, Frequency of reads aligned to *Apoa1* and off-target (OT) sites (including or not ITR sequences) relative to reads aligned to the mouse genome. **d**, Count of ITR-containing reads mapping to off-target sites per mouse. **e**, Count of unique off-target integration sites per mouse. Start mapping positions were consolidated using a 1000-bp sliding window for unique off-target site identification. **f**, Distribution of unique off-target integration sites per chromosome. **g**, Distribution of unique off-target integration sites in intergenic regions, introns, promoters - transcriptional start sites (TSS), exons and translational termination sites (TTS). **h**, Redundant off-target sites found in Unselected mice. **i**, Redundant off-target sites found in Repair Drive mice. In **h** and **i**, the read count is shown for each site. Sites are shown in descending order of redundancy and read count. Sites identified in both Unselected and Repair Drive mice are indicated with an asterisk (*Gm10800*, *Cps1*, *Ass1* and *Alb*). Data are expressed as mean ± s.d. (n= 5 Unselected and Repair Drive mice) with significance determined by two-tailed student’s t-test. * p<0.05, ** p<0.01, *** p<0.001 and **** p<0.0001.

### Repair Drive enables safe and durable expression of a therapeutic transgene

To address the long-term transgene expression and safety of Repair Drive, we injected adult mice with an AAV-Donor to integrate the therapeutic transgene Factor IX (*FIX*) – deficient in patients with Hemophilia B - into the *Apoa1* locus. We selected targeted hepatocytes with *Fah*-siRNA for twelve weeks and euthanized the mice after one year (Fig. 6a). Repair Drive resulted in enrichment of *Apoa1* alleles with HDR integration of the AAV-Donor (Extended Fig. 4a). FIX secretion increased ∼5-fold in Repair Drive mice reaching therapeutic levels and was sustained over time-342.04±162.18 vs. 74.14±26.64 ng/ml in Unselected mice at endpoint (Fig. 6b). Hepatocyte cell death and proliferation were not elevated in Repair Drive mice at endpoint (Extended Fig. 4c and Supplementary Fig. 23). Several mice had liver steatosis, which was expected in aged *C57BL6/J* mice and was not associated with treatment group (Supplementary Fig. 24). Careful examination of all livers for potential abnormalities^24^ revealed one hemangioma in a Control liver, and localized proliferative lesions in four livers-two in Unselected and two in Repair Drive mice (Supplementary Fig. 25-26 and Table S9). Most importantly, the livers of Repair Drive mice had large colonies of targeted hepatocytes, normal histology, and fully restored Fah protein expression after a year of follow up (Fig. 6c).

**Fig. 6.**
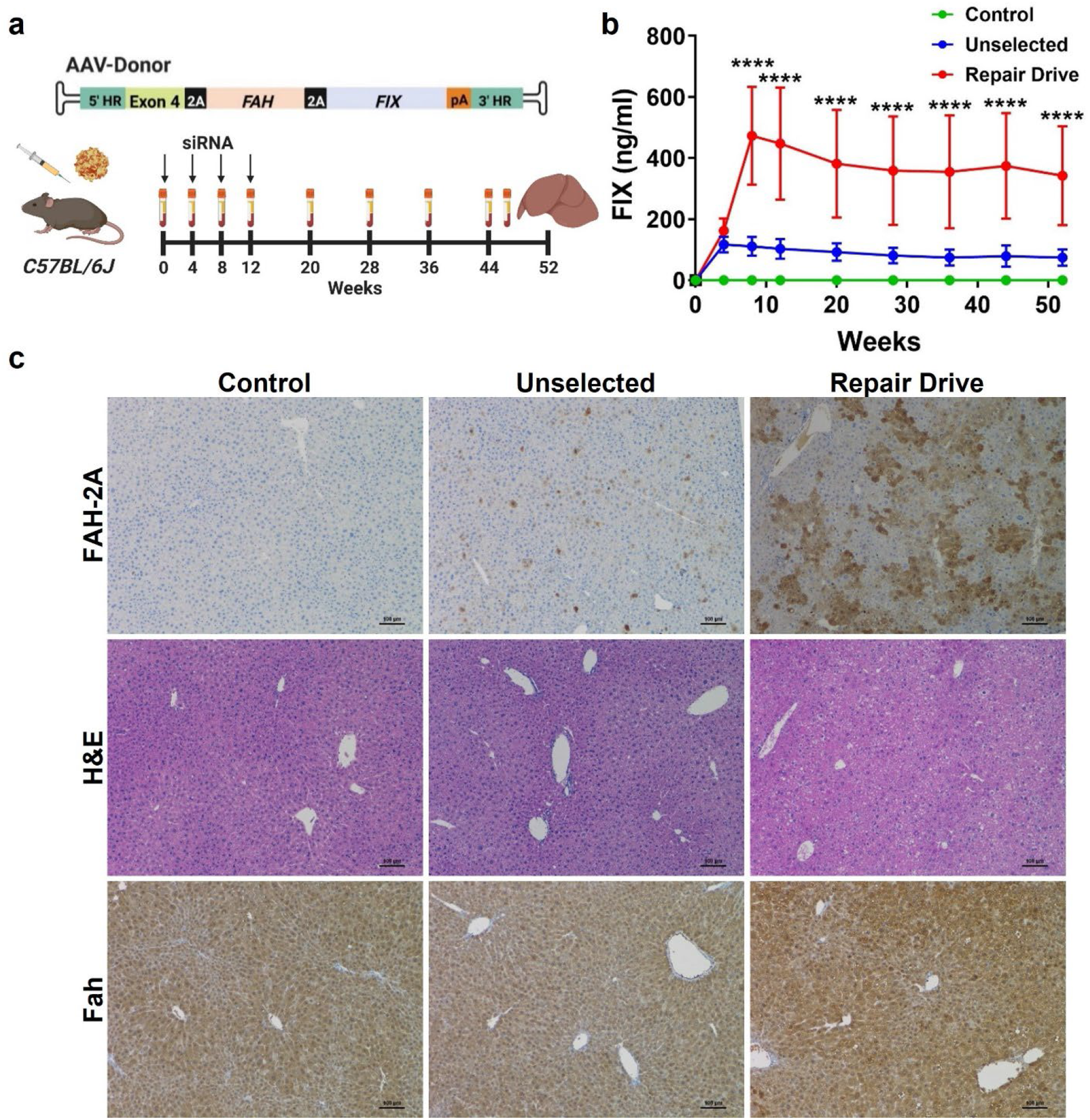
Long-term transgene expression and safety following Repair Drive. **a**, Eight-week-old *C57BL/6J* mice were injected with 5×10^11^ GC of AAV-CRISPR and 1×10^11^ GC of an AAV-Donor containing the promoterless coding sequences of human *FAH* (selectable marker) and *FIX* or 6×10^11^ GC of AAV-GFP as control. Mice were injected with *Fah*-siRNA (3 mg/kg or saline as control) every 4 weeks until 12 weeks post-AAV injection. Blood was collected at time 0 and every two to four weeks and mice were sacrificed at 52 weeks post AAV-injection. Created with BioRender.com. **b**, Human FIX ELISA in plasma samples. **c**, Representative immunohistochemistry staining of 2A-tagged FAH (FAH-2A), hematoxylin and eosin (H&E) and Fah in the liver. Scale bar is 100 µm. Data are expressed as mean ± s.d. (n= 3 Control and 9 Unselected and Repair Drive mice) with significance determined by two-way ANOVA followed by Tukey test. **** p<0.0001 Repair Drive vs. Control and Unselected mice.

## Discussion

In this study, we demonstrate the potential of Repair Drive, a novel strategy for selectively expanding gene-targeted hepatocytes *in vivo*, that can reach ∼25% of all hepatocytes. As a result, the expression of the therapeutic transgene *FIX* increased 5-fold, reaching therapeutic levels. Despite transient liver damage, Repair Drive was safe and well tolerated in the long-term and did not increase hepatocellular carcinoma (HCC) risk in ∼14-month-old mice. Comprehensive analysis of editing events revealed that Repair Drive resulted in significant enrichment for HDR-targeted *Apoa1* alleles over unwanted on- and off-target editing events.

In our previous work, we identified *Apoa1* as an effective integration site for liver-directed gene therapy. We showed that this locus enabled the sustained expression of therapeutic proteins, ameliorating hypercholesterolemia in neonatal apolipoprotein E (*Apoe*) knockout mice and correcting hereditary tyrosinemia type I (HT-1) in *Fah* knockout mice.^17^ Despite its effectiveness in neonatal mice, gene targeting efficiency remains low in the adult liver. Similarly, in recent work the *Alb*-targeted expression of *FIX* efficiently corrected Hemophilia B in neonatal mice, yet it was inadequate in adult mice.^25^ To increase the efficiency of gene targeting at the *Alb* locus, Tsuji et al administered ribonucleotide reductase inhibitors to neonatal and juvenile mice resulting in enhanced targeted integration.^8^ However, the effectiveness of this strategy still needs to be evaluated in the adult liver, where HDR is rare. Other approaches conferred a selective growth advantage to gene-modified hepatocytes through permanent inactivation of important metabolic genes-*Hpd* and *Cypor*.^11, 14, 15^ Although they effectively expanded edited cells increasing transgene expression, these strategies have safety concerns.^26–31^ Sustained knockdown of *Hpd* could mimic hereditary tyrosinemia type III, and potentially cause neurotoxic levels of tyrosine.^26, 27^ Likewise, the permanent removal of *Cypor* may adversely impact the homeostasis of lipids and bile acids and the metabolism of hormones and drugs.^28–31^ In fact, deletion of this gene leads to a growth disadvantage under homeostatic conditions^32^ and severe steatohepatitis.^28, 30^

Repair Drive has several advantages over those approaches: (1) it increases precise gene insertion in the adult liver by expanding correctly HDR-targeted cells; (2) it involves transient inhibition of an essential gene, whose expression is fully restored following expansion; (3) targeted cells do not contain permanent disruption of other genes; (4) the selection pressure can be adjusted. It can be raised by increasing the siRNA dose or by combining the siRNA with a high-protein diet. Otherwise, it can be reduced by siRNA removal or dose reduction. In principle, Repair Drive should also be achievable using other essential genes for selection, provided potent inhibition can be achieved with a small molecule or siRNA, and their expression can be effectively coupled to the gene-editing event.

One major concern with CRISPR/Cas9 is the high level of unintended editing events in the on-target site. In addition to indel formation, we and others reported NHEJ-mediated insertion of AAV genomes at the cut site including ITRs.^17, 33–35^ Here, we used droplet digital PCR (ddPCR) and long-read sequencing to comprehensively characterize gene editing events at the *Apoa1* locus. We found that Repair Drive resulted in strong enrichment of HDR-modified *Apoa1* alleles – up to 6-fold by ddPCR. Interestingly, we observed heterogeneous HDR events. In addition to correct HDR, we detected alleles with HDR integration at the 5’ end and ITR insertion at the 3’ end, and/or with partial truncation of TdTomato. Their enrichment following expansion suggests that they all resulted in functional expression of FAH. TdTomato truncation was most likely due to recombination within this repetitive transgene. In parallel, we observed significant decreases of NHEJ-derived indels formation and AAV genome insertion. Despite this, the rate of unwanted editing events did not go to zero. Indeed, these events may still persist as passengers in cells harboring an HDR-modified *Apoa1* allele. In addition, we observed evidence of AAV-CRISPR integration, which could result in permanent expression of Cas9. Repair drive should also be compatible with other delivery methods, such as transient delivery of Cas9 mRNA,^1, 36, 37^ to further reduce the likelihood of unintended integration events observed with AAV.

Off-target integration of AAV is another major concern in AAV gene therapy. Although mostly episomal, AAVs can integrate into the host genome with a frequency of 0.1 to 3%^38, 39^ and potentially result in tumorigenesis.^40, 41^ Indeed, AAVs were found integrated in the RNA imprinted and accumulated in nucleus (*Rian*) locus in AAV-injected neonatal mice, which led to HCC development.^22, 42^ Here, we performed a GISA-seq assay for unbiased genome-wide retrieval of AAV-Donor integration. Our data showed up to ∼230 unique integration sites per mouse. However, this could be underestimated by several technical limitations. First, most of the reads mapped to episomal AAVs, which can potentially interfere with the detection of rare integration events. Second, our GISA-seq restricted the search to the 3’ end of AAV-Donor, therefore missing the integration of other portions of the AAV-Donor as well as AAV-CRISPR. Finally, we potentially missed certain integration outcomes in highly repetitive genomic regions. Nevertheless, we observed recurrent insertion in previously reported AAV integration hotspots – *Alb*, *Afp*, *Ass* and *Gm10800*.^22, 23^ Most importantly, the frequency of off-target integration dramatically decreased following Repair Drive, suggesting that they did not result in a favorable transgene expression from AAV-Donor. On the contrary, the reads mapping to *Apoa1* strongly increased. However, we also observed enrichment of integration events within highly transcribed loci – *Alb*, *Afp*, *Cps1*, *Ass1* - which could potentially contribute to a residual off-target expression of the Donor cassette. Another important finding is that we did not detect insertional mutagenesis in proto-oncogenes, including the *Rian* locus. In line with these data, we observed no HCC following expansion, at least within the timeframe of these experiments (up to one year).

Overall, we show that Repair Drive enables the selective expansion of gene-targeted hepatocytes *in vivo*. Repair Drive could in principle be amenable to use with other gene editors, editing approaches, and delivery systems. This approach should broaden the range of liver diseases correctable with gene therapy using targeted transgene insertion or HDR.

## Methods

### Plasmid design and cloning

The AAV-CRISPR plasmid (1507-pAAV-U6-Apoa1-gRNA2-SA-HLP-SaCas9-HA-OLLAS-spA) was generated as previously described.^17, 43^ This plasmid encodes the gRNA for targeting the 3’ UTR of *Apoa1* (GTTTATTGTAAGAAAGCCAATGC)^17^ under the control of a U6 promoter, and *Staphylococcus aureus* Cas9 (SaCas9) under the control of the hybrid liver-specific promoter (HLP). The AAV donor plasmids were generated by standard molecular biology approaches starting from 1731-pAAV-Apoa1-Target-2A-FIX-pA and 1771-pAAV-Apoa1-Target-2A-FAH-pA plasmids.^17^ The donor plasmids include exon 4 of the murine *Apoa1* gene (NC_000075.6), but introduce the promoterless transgene coding sequences in place of the endogenous stop codon, flanked by homology arms (HR) to the *Apoa1* locus on each side (5’HR: 700 bp; 3’HR: 480 bp).^17^ In place of the stop codon, each donor plasmid includes a Glycine-Proline-Glycine-P2A sequence (referred as 2A) in frame with the human *FAH* coding sequence (without stop codon), followed by another 2A sequence. Then, the TdTomato or human *FIX* coding sequence was cloned between the second 2A sequence and a small synthetic poly(A) signal, resulting respectively in 1948-pAAV-ApoA1-Target-2A-hFAH-2A-Tom-pA and 1926-pAAV-ApoA1-Target-2A-hFAH-2A-hFIX-pA.

### AAV production

Recombinant AAV8 vectors were generated as previously described with several modifications.^17, 44^ Plasmids required for AAV packaging, adenoviral helper plasmid pAdDeltaF6 (PL-F-PVADF6) and AAV8 packaging plasmid pAAV2/8 (PL-T-PV0007) were obtained from the University of Pennsylvania Vector Core. Each AAV transgene construct was co-transfected with the packaging constructs into 293T cells (ATCC, CRL-3216) using polyethylenimine (PEI). Cell pellets were harvested and purified using a single cesium chloride density gradient centrifugation. Fractions containing AAV vector genomes were pooled and then dialyzed against PBS using a 100kD Spectra-Por® Float-A-Lyzer® G2 dialysis device (Spectrum Labs, G235059) to remove the cesium chloride. Purified AAV were concentrated using a Sartorius™ Vivaspin™ Turbo 4 Ultrafiltration Unit (VS04T42) and stored at -80°C until use. AAV titers were calculated after DNase digestion using qPCR relative to a standard curve of plasmids containing the appropriate transgene portion. Primers used for titer are included in Table S10.

### *In vitro* screening of *Fah*-siRNAs

Primary mouse hepatocytes were purchased from BioIVT and cultured in INVITROGRO medium with 10% of fetal bovine serum. Hep3b cells were purchased from ATCC and cultured in Gibco EMEM with 10% fetal bovine serum. Cells were transfected with 0.1, 1, 10 and 50 nM of 22 *Fah*-siRNAs in 384-well format using Lipofectamine RNAi Max for 24 hours, following the manufacturer’s instructions. RNA was extracted using an automated protocol on a BioTek-EL406 platform using the Dynabeads™ mRNA DIRECT™ Purification Kit (Invitrogen™, Catalog No. 61012). cDNA was synthesized using the High-Capacity cDNA Reverse Transcription Kit with RNase Inhibitor (Applied Biosystems™, Catalog No. 4374967) according to the manufacturer’s recommendations. A master mix containing 1 µL 10X Buffer, 0.4 µL 25X deoxyribonucleotide triphosphate, 1 µL 10X Random primers, 0.5 µL Reverse Transcriptase, 0.5 µL RNase inhibitor, and 6.6 µL of H_2_O per reaction was added to RNA isolated above. mRNA levels were quantified by performing RT-qPCR analysis. 2 μL of cDNA were added to a RT-qPCR master mix containing 0.5 μL of 20X human or mouse GAPDH TaqMan probe, 0.5 μL 20X FAH probe (ThermoFisher Scientific Hs001646611_m1 [human] and Mm00487336 [mouse]) and 5 μL LightCycler 480 Probes Master mix per well in 384 well plates. The RT-qPCR assay was carried out in a LightCycler480 Real Time PCR system (Roche) using the TaqMan gene expression assay. To calculate relative fold change, real-time data were analyzed using the Delta-Delta Threshold Cycle (Relative Quantification) (ΔΔCt[RQ]) method^45^ and normalized to control assays performed using cells transfected with 10 nmol/L non targeting siRNA control.

### Animals

*C57BL/6J* mice (stock number: 000664) were obtained from Jackson Laboratories. Animals were allowed free access to food and water and maintained on a standard chow diet. Where indicated, mice were fed with a high-protein diet (70% casein, TD.03637, Envigo). AAV-CRISPR (5 × 10^11^ genome copies (GC)) and AAV-Donor (1 × 10^11^ GC) were diluted in 300 µl of sterile saline and intraperitoneally injected to 8-week-old male mice. Control mice were injected with 6 × 10^11^ GC of AAV8-CB-EGFP (AAV-GFP). All treatment conditions were randomly allocated within each cage of mice at the time of AAV injection. In addition, mice were subcutaneously injected with 3 mg/kg of *Fah*-siRNAs or saline as control. Mice were fasted 5 hours prior to injection and before subsequent blood collection. Blood was collected via retro-orbital bleeding using heparinized Natelson collection tubes, and plasma was isolated by centrifugation at 10,000 g for 20 minutes at 4°C. For tumor analysis, livers were examined for abnormalities by “breadloaf” sectioning at 3-mm intervals as previously described.^24^ Raised nodular lesions and discolored areas (suspected tumors) were collected and fixed in 10% formalin for histology analysis. All experiments were approved by the Baylor College of Medicine Institutional Animal Care and Use Committee (IACUC) and performed in accordance with institutional guidelines under protocol number AN-7243.

### Inference of CRISPR Edits (ICE) analysis

ICE analysis of indel formation at the *Apoa1* locus was performed as previously described.^17^ Primer sequences used for amplifying the *Apoa1* locus are listed in Table S10.

### Integration PCR

Genomic DNA was extracted from livers using the DNeasy Blood and Tissue kit (Qiagen) following the manufacturer’s protocol. Integration PCR was performed as previously described with some modifications.^17^ Briefly, the *Apoa1* locus was PCR-amplified to detect integration of the AAV-Donor by using a forward primer within an endogenous genomic region upstream of the 5’HR and the reverse primer within the *FAH* coding sequence. APEX TaqRed Master Mix (Apex Bioresearch Products) was used following the manufacturer’s protocol and the PCR products were separated by agarose gel electrophoresis. Primers sequences are available in Table S10.

### Droplet digital PCR (ddPCR)

ddPCRs for quantifying the AAV-Donor homology-directed repair (HDR)- and non-homologous end joining (NHEJ)-mediated integration events at the *Apoa1* locus were performed as previously described.^17^ An EvaGreen-based ddPCR assay was used for quantifying the vector copy number (VCN) for AAV-CRISPR and AAV-Donor. Primer pairs targeting SaCas9, *FAH* and Transferrin receptor (*Tfrc*) were respectively used for amplifying AAV-CRISPR, AAV-Donor and *Tfrc* as reference gene. The reaction mixes were prepared using 15 ng of genomic DNA, QX200™ ddPCR™ EvaGreen Supermix (Bio-Rad), 200 nM target primers and 10 U of Hind III–HF restriction enzyme (NEB) in a final volume of 20 μL. PCR was performed according to the manufacturer’s cycling protocol. VCN for AAV-CRISPR and AAV-Donor were normalized to *Tfrc* to determine genome copies per diploid cells. Primer and probe sequences used for ddPCR are listed in Table S10.

### On-target gene editing analysis

Genomic DNA was extracted from the liver of one Control, five CRISPR/Donor (Unselected) and five CRISPR/Donor + siRNA (Repair Drive) mice. A PCR reaction with two amplification cycles (PCR1) was used to amplify 1-6 kb genomic regions at the *Apoa1* target site and simultaneously tag the 5′ and 3′ ends of each template molecule with 18 bp terminal Unique Molecular Identifier (UMI) using a tailed primer pair. The first section of both tailed primers was a synthetic priming site used for downstream amplification, followed by the 18 nucleotides “patterned” UMI (NNNYRNNNYRNNNYRNNN) and target-specific sequences. After UMI labeling, each strand of the DNA duplex was tagged with a unique combination of dual UMI. A second PCR (PCR2) was used to amplify the UMI-tagged DNA molecules. A third PCR (PCR3) was used to reamplify the PCR2 product for generating barcoded amplicons for multiplexed sequencing. The final barcoded PCR3 amplicon products contained the symmetric barcode sequences at both ends. Barcoded amplicon samples from PCR3 were pooled and used for SMRT-seq library preparation (SMRTbell® prep kit 3.0, Pacific Biosciences). The SMRTbell library was sequenced on a PacBio Sequel II 8M flow cell in CCS mode at the Human Genome Sequencing Center (Baylor College of Medicine), and HiFi reads were produced. HiFi reads with an average quality score of Q40 (99.99%) single-molecule read accuracy were demultiplexed and processed by the longread_umi pipeline^46^ to generate UMI consensus reads. UMI consensus sequences were aligned to the reference amplicon sequence using Burrows-Wheeler Alignment (BWA)-Maximal Exact Match (MEM).^47^ We used a custom Matlab script to analyze the UMI consensus sequences and characterize the gene editing outcomes. In short, gene modification variants were categorized into ten groups: (a) unmodified, (b) large deletions, (c) small deletions, (d) small insertions, (e) HDR, (f) HDR with truncation of one copy of TdTomato – HDR(-Td), (g) 5’HDR-3’NHEJ, (h) 5’HDR(-Td)-3’NHEJ, (i) AAV capture with ITR at the integration junction and (j) ITR-less AAV capture. Primer sequences are available in Table S10.

### Genome-wide analysis of AAV-Donor off-target integration sites

We developed a <u>G</u>enome-wide Integration <u>S</u>ite of <u>A</u>AV-Donor identified by <u>seq</u>uencing (GISA-seq) to determine the integration sites of AAV-Donor in the mouse genome. GISA-seq is a modified version of an anchored PCR reaction described in the GUIDE-seq protocol.^48^ A single primer is designed to anneal to TdTomato sequence in the AAV-Donor and amplify outward to sequence the regions across junctions. Genomic DNA was extracted from the liver of one Control, five CRISPR/Donor (Unselected) and five CRISPR/Donor + siRNA (Repair Drive) mice. 4 µg of DNA in 80 µl reaction were incubated at 37°C for seven minutes and randomly fragmented to 300 bp – 4 kb size products using NEBNext® dsDNA Fragmentase (New England BioLabs) and then cleaned using 0.5x SPRI beads. After end-repair and A-tailing, unique Y-adapters containing UMI sequences and sample-specific P5 indexes were ligated to the DNA using NEBNext® Ultra™ II FS DNA Library Prep Kit for Illumina (New England BioLabs). Y-adapter-ligated DNA fragments were amplified by PCR1 using a TdTomato-specific forward primer and a reverse primer binding to the P5 sequence on Y-adapter. Subsequent nested PCR2 was performed to enrich the target amplicons and improve specificity. Nested PCR3 was performed using staggered forward primer annealing at 3’HR and sample-specific P5 index reverse primer to add adapter sequences and reduce amplicon size. PCR4 was performed using index primers to add sample-specific P7 index. Pooled library was sequenced using MiSeq Reagent Kit v3 (600-cycle) (Illumina). Selected integration sites were validated by PCR amplification and Sanger sequencing. Primer and adapter sequences are available in Table S10.

### Pipeline design of GISA-seq

Reads that share the same UMI and the same first six bases of genomic sequence are presumed to originate from the same pre-PCR molecule and are thus consolidated into a single consensus read to improve quantitative interpretation of GISA-seq read counts. Sample demultiplexing and PCR duplicate consolidation steps followed GUIDE-seq pipeline.^48^ The demultiplexed, consolidated Read2 was processed using an ad-hoc bioinformatics pipeline. A custom reference genome was built including mm10, ITR-less AAV-CRISPR, ITR-less AAV-Donor and ITR sequences. Reads generated from non-specific amplification were filtered using a target-specific 8 bp sequence. Next-generation sequencing (NGS) sequences were aligned against the custom reference genome using BWA-MEM, and reads unable to map properly to the reference were removed during the filtering step. The resulting Read2 contained a 53 bp long 3’HR region followed by the sequence of interest. Multiple filters were applied to remove episomal AAV sequences: (1) AAV-CRISPR filter removed reads mapping to AAV-CRISPR and not to mm10; (2) AAV-Donor filter removed reads mapping to AAV-Donor and excluding the 53 bp long 3’HR region; (3) Complex recombination of AAV-Donor and AAV-CRISPR were detected and removed. The resulting reads had AAV-Donor integration in mm10 and were further categorized into (4) on-target (ON) AAV-Donor integration at the *Apoa1* locus or (5) off-target (OT) AAV-Donor integration. OT AAV-Donor integration reads were further categorized based on the presence or absence of ITR at the integration junction. For ONs, start mapping positions were consolidated using a 100-bp sliding window. For OTs, start mapping positions were consolidated using a 1000-bp sliding window. The sequence was retrieved from each OT integration site and re-aligned to the mm10 using a BWA-MEM to generate BED (Browser Extensible Data) files using SAMtools^49^ and BEDTools^50^ for further annotation using HOMER (Hypergeometric Optimization of Motif EnRichment) suite.^51^ Identified AAV-Donor integration site from each mouse, sorted by GISA-seq read count were annotated in a final output table (Table S7 and S8).

### Data and materials availability

All data needed to evaluate the conclusions in the paper are present in the paper and/or the Supplementary Materials. PacBio and Illumina sequencing data are accessible at the Sequence Read Archive (SRA) under accession PRJNA954356. Source code and analysis scripts for GISA-seq are available on GitHub at baolab-rice · GitHub.

### Immunohistochemistry

Livers were fixed in 10% formalin overnight at 4°C, then gradually dehydrated with ethanol and embedded in paraffin. Immunohistochemistry and hematoxylin and eosin (H&E) staining were performed by the Texas Digestive Diseases Morphology Core at Baylor College of Medicine, as previously described.^17^ The following primary antibodies were used for immunohistochemistry: rabbit anti 2A peptide (1:7500, ABS31, Sigma-Aldrich), rabbit anti Ki67 (1:60, CRM325, Biocare); rabbit anti FAH (1:65, ab151998, Abcam); rabbit anti Cd34 (1:250, ab81289, Abcam). The Ki67, FAH and Cd34 antibodies were then detected with a Rabbit-on-Rodent HRP-Polymer (RMR622H, Biocare) and visualized with DAB chromogen (DB801, Biocare). The 2A antibody was detected and visualized using a Leica Bond Polymer Detection kit (DS9800). Tunel staining was performed and detected using ApopTag Peroxidase In Situ Apoptosis Detection Kit (Millipore, S7100). Reticulin staining was performed using the Epredia Reticulin Sliver Stain Kit (87025, Epredia), following the manufacturer instructions. Slides were counterstained with hematoxylin, dehydrated, and mounted with a permanent mounting medium. A Nikon Ci-L bright field microscope was used for imaging at the Integrated Microscopy core (Baylor College of Medicine).

### Direct immunofluorescence

Livers were fixed in 4% PFA (PBS) for 24 hours at 4°C, then incubated in 30% sucrose (PBS) for 24 hours at 4°C and finally embedded in OCT medium. Frozen tissue blocks were sectioned at 14 µm and stained with DAPI before mounting using Prolong Diamond (Invitrogen). Sections were imaged using Zeiss Axio Scan.Z1 scanner at the RNA In Situ Hybridization core (Baylor College of Medicine) with a 20x 0.8 lens and appropriate filters and analyzed using ZEN 3.3 software (ZEN lite). TdTomato-positive cells quantification was performed by manual count of positive hepatocytes in three 1 mm^2^ fields per liver across the whole section. Nuclei were quantified using ImageJ.

### Western blot

Liver protein isolation and western blots on liver extracts and plasma samples were performed as previously described.^17^ Primary antibodies to the 2A peptide (1:5000, rabbit, ABS31, Sigma-Aldrich), GFP (1:3000, rabbit, A-11122, Fisher-Scientific), FAH (1:1000, rabbit, SAB2100745, Sigma-Aldrich), apoA1 (1:5000, rabbit, K23500R, Meridian) and beta tubulin (1:500, mouse, University of Iowa Developmental Studies Hybridoma Bank E7) were diluted in 1% BSA in PBS-T and membranes were incubated overnight at 4°C. Goat anti-rabbit 680 nm and anti-mouse 800 nm antibodies (1:15000, 611-144-002-0.5 and 610-145-002-0.5, Rockland) were incubated at room temperature for 1 hour and imaged using an Odyssey Classic (Li-Cor). Red Ponceau stain was used as loading control for western blot on plasma samples.

### Plasma analysis

Alanine aminotransferase (ALT) was measured using the Teco ALT (SGPT) Kinetic Liquid Kit (A524-150), following the manufacturer’s protocol. Human FIX was measured in 1:16.7 diluted plasma by using Factor IX Human SimpleStep ELISA Kit (ab188393, Abcam), following the manufacturer’s protocol.

### Nuclei isolation from liver and library generation

∼ 100 mg of frozen liver were used for nuclei isolation and single nucleus RNA sequencing (snRNAseq) analysis. Nuclei were isolated from one Control and one CRISPR/Donor + siRNA (Repair Drive) mouse sacrificed at 12 weeks post-AAV injection. Liver samples were cut into tiny pieces in 250 µL of buffer HB (0.25 M Sucrose, 25 mM KCl, 5 mM MgCl_2_, 20 mM tricine-KOH, pH 7.8) supplemented with 1x DTT, spermine and spermidine, and 0.2 U/µL of Protector RNase inhibitor (Millipore Sigma) (referred as buffer HB++). Then, samples were centrifuge at 500 g for 5 minutes and resuspended in 500 µL of buffer HB++. Samples were homogenized in a Dounce homogenizer firstly with a loose pestle for 30 strokes, followed by a tight pestle for 10 strokes after adding 250 µL of 1% NP-40 (in buffer HB++). Homogenates were filtered through a pre-wet 40 µm cell strainer (Falcon) by adding 10 mL of nuclei wash buffer (NWB, 1% BSA in PBS supplemented with 0.2 U/µL of Protector RNase inhibitor). Samples were centrifuged at 500 g for 10 minutes at 4°C using a swinging bucket rotor and the pellets were resuspended in 500 µL of NWB buffer and filtered through a 40 µm cell strainer (Flowmi, Millipore Sigma). 900 µL of sucrose cushion buffer (SCB: 9 volumes of Nuclei PURE 2M sucrose cushion solution and 1 volume of Nuclei PURE sucrose cushion solution, Sigma-Aldrich, supplemented with 0.2 U/µL of Protector RNase inhibitor) were added to the resuspended pellet and the obtained mix was layered on top of 500 µL of SCB in a 2 mL Eppendorf tube. Nuclei were centrifuged at 13000 g for 45 minutes at 4°C. The obtained pellet was resuspended in 1 mL of NWB and filtered through a pre-wet 20 µm CellTrics filter (Fisher Scientific). Nuclei were finally centrifuged at 500 g for 5 minutes and resuspended in 200 µL of NWB buffer. Nuclei were stained with DAPI and nuclei purity and integrity were confirmed using a fluorescent microscope. Samples were diluted to obtain 15000 - 18000 nuclei for single cell library construction according to the 10X single cell gene expression workflow (10X Genomics, Chromium Next GEM Single Cell 3’ Kit v3.1, 1000269). Quality of the cDNA and the libraries were analyzed using the high sensitivity NGS fragment analysis kit (Agilent). Libraries were sequenced on an Illumina NovaSeq 6000 instrument.

### SnRNAseq analysis

Raw reads were aligned to a custom mm10 reference file that included TdTomato and GFP reference sequences. CellRanger Count estimated 17117 nuclei from the combined Control and Repair Drive samples, averaging 30K reads per nucleus with a median of 2000 unique genes per nucleus. Ambient background RNA was removed after alignment using CellBender.^52^ Single nuclear RNA-seq data was analyzed using the latest Seurat v4.0 package suite.^53^ Quality control filters were used to remove low quality nuclei. All genes that were not expressed in at least 10 nuclei were excluded, and nuclei with less than 200 genes were removed. 15979 nuclei and 18153 genes remained after filtration. Nuclei Doublets were determined both manually and through DoubletFinder.^54^ RNA count was normalized using SCT from Seurat V4 (SCTransform) regressing out any contaminating mitochondria mRNA. Principle components analysis (PCA) was determined by RunPCA and both samples were integrated using Harmony package.^55^ The number of PCA to be used for downstream analysis was determined using elbowplot and jackstraw for downstream analysis. We used Uniform Manifold Approximation and Projection (UMAP) for dimensional reduction and visualization of the single nuclear data. For determining hepatocyte zonation and ploidy prediction, we used a set of previously identified markers.^19, 20^ SnRNAseq datasets are reported in Table S1-S4.

### Statistics

Graphpad Prism 9 was used for statistical analyses. All data are shown as the mean +/- standard deviation. Comparisons involving two groups were evaluated by a two-tailed student’s t-test. For comparisons involving three or more groups, a one-way or two-way ANOVA was applied, with Tukey post-test used to test for significant differences among groups. In all cases, significance was assigned at p<0.05.

## Acknowledgements

This work was supported by US National Institute of Health grants U42OD026645, DK124477, HL132840 to W.R.L., UG3 HL151545 to G.B and W.R.L., R01HL152314 to G.B., DK115461 to K.D.B, and American Heart Association 19POST34430092 to M.D.G. This work was also supported by the Texas Digestive Diseases Morphology Core (P30DK56338) and the Integrated Microscopy Core (DK56338 and CA125123), and with assistance from the Baylor College of Medicine Gene Vector Core.

## Contributions

M.D.G., A.H., M.A.C., C.W., A.M.D., M.N.F. performed *in vivo* experiments. S.H.P., M.C., L.S. performed sequencing and bioinformatics analysis of genome editing. A.C., S.H., T.C., S.L., K.W., P.M., J.Q., M.K.S., I.Z. and V.J. designed, tested and supplied siRNA reagents, M.C.L. performed sectioning, histology and imaging, K.R.P. assessed pathology of liver biopsies, R.G.L., J.K., J.F.M. performed and interpreted single nucleus RNA-Seq, M.D.G., S.H.P., A.C., K.D.B., J.F.M., V.J., G.B., and W.R.L. provided critical intellectual input to the project. W.R.L. conceptualized the Repair Drive strategy. M.D.G., S.H.P., A.C., V.J., G.B., and W.R.L. drafted the manuscript which was edited and approved by all authors.

---

**Extended Data Fig. 1.**
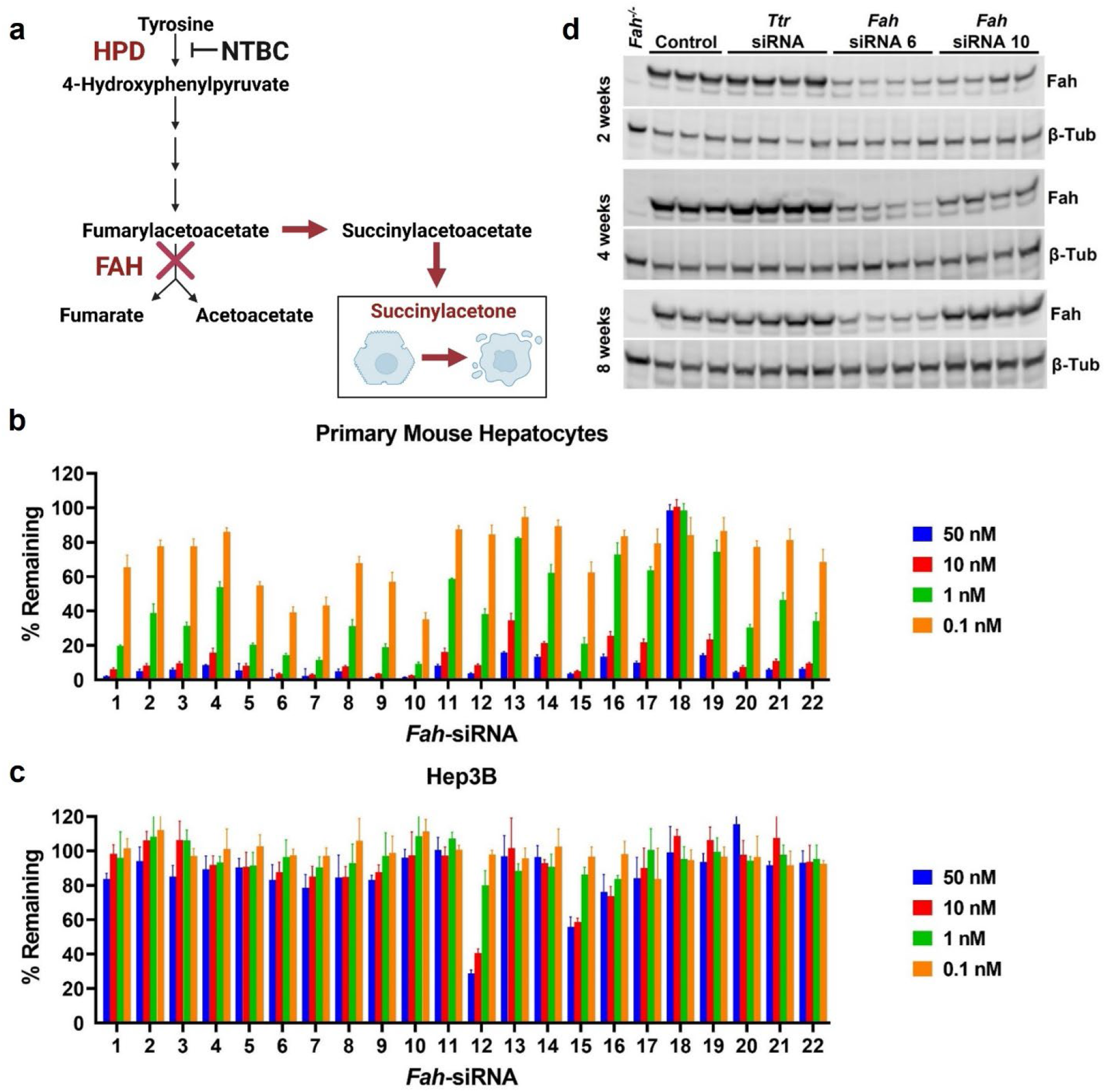
Fah knockdown. **a**, Simplified scheme of the tyrosine catabolic pathway. Mutations or knockdown of Fah cause accumulation of upstream hepatotoxic metabolites - succinylacetone. Liver injury can be prevented by blocking the pathway upstream through Nitisinone (NTBC)-mediated inhibition of Hpd. Created with BioRender.com. **b** and **c**, *In vitro* screening of twenty-two *Fah*-siRNA duplexes. **b**, qPCR analysis of murine *Fah* expression in primary mouse hepatocytes incubated with 0.1, 1, 10 and 50 nM of *Fah*-siRNAs for 24 hours. **c**, qPCR analysis of human *FAH* expression in Hep3B cells transfected with 0.1, 1, 10 and 50 nM of *Fah*-siRNAs for 24 hours. Data are shown as percentage of remaining mRNA relative to untreated cells. Data are expressed as mean ± standard deviation (n=4 replicates). **d**, Western blot analysis of Fah knockdown in liver lysates from mice injected with 3 mg/kg of two different *Fah* siRNAs at 2-, 4- and 8-weeks post-injection. A siRNA for transthyretin (*Ttr*) was used as delivery control. Liver lysates from Fah knockout (*Fah*^-/-^) mice were used as negative control for the antibody. Beta

**Extended Data Fig. 2.**
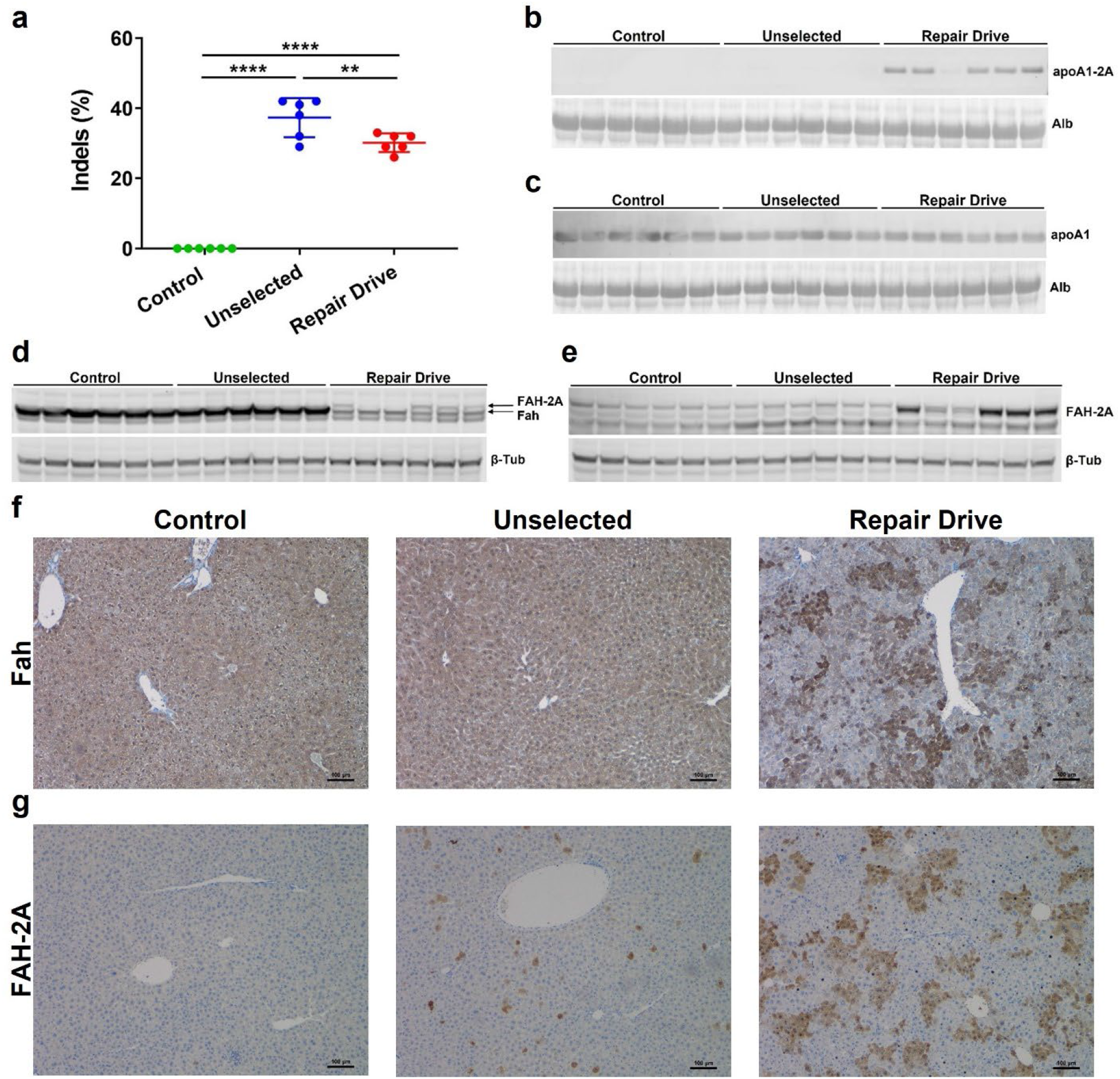
Enrichment of transgenic products following Repair Drive. **a**, Frequency of indel formation at the *Apoa1* 3’ UTR by Inference of CRISPR Edits (ICE) analysis. Western blot analysis of 2A-tagged (**b**) and total apoA1 (**c**) in plasma with Alb as loading control. Western blot analysis of endogenous Fah (**d**) and FAH-2A (**e**) in liver lysates with β-Tub as loading control. Representative immunohistochemistry staining of Fah (**f**) and FAH-2A (**g**) in the liver. Scale bar is 100 µm. Data are expressed as mean ± s.d. (n= 6 mice per group), with significance determined by one-way ANOVA followed by Tukey test. ** p<0.01 and **** p<0.0001.

**Extended Data Fig. 3.**
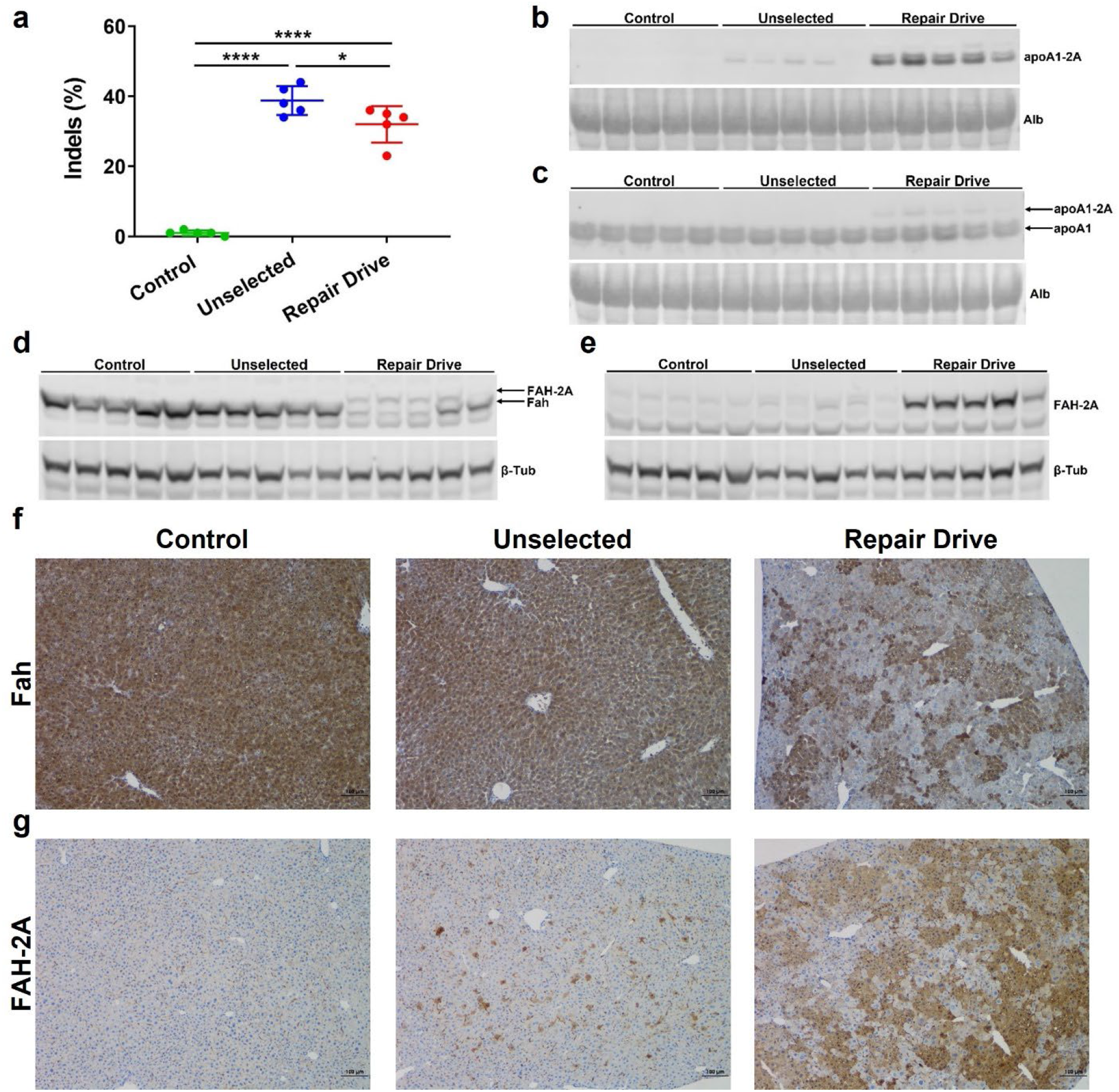
Enrichment of transgenic products following Repair Drive with high protein diet. **a**, Frequency of indel formation at the *Apoa1* 3’ UTR by ICE analysis. Western blot analysis of 2A-tagged (**b**) and total apoA1 (**c**) in plasma with Alb as loading control. Western blot analysis of endogenous Fah (**d**) and FAH-2A (**e**) in liver lysates with β-Tub as loading control. Representative immunohistochemistry staining of Fah (**f**) and FAH-2A (**g**) in the liver. Scale bar is 100 µm. Data are expressed as mean ± s. d. (n= 5 mice per group), with significance determined by one-way ANOVA followed by Tukey test. * p<0.05 and **** p<0.0001.

**Extended Data Fig. 4.**
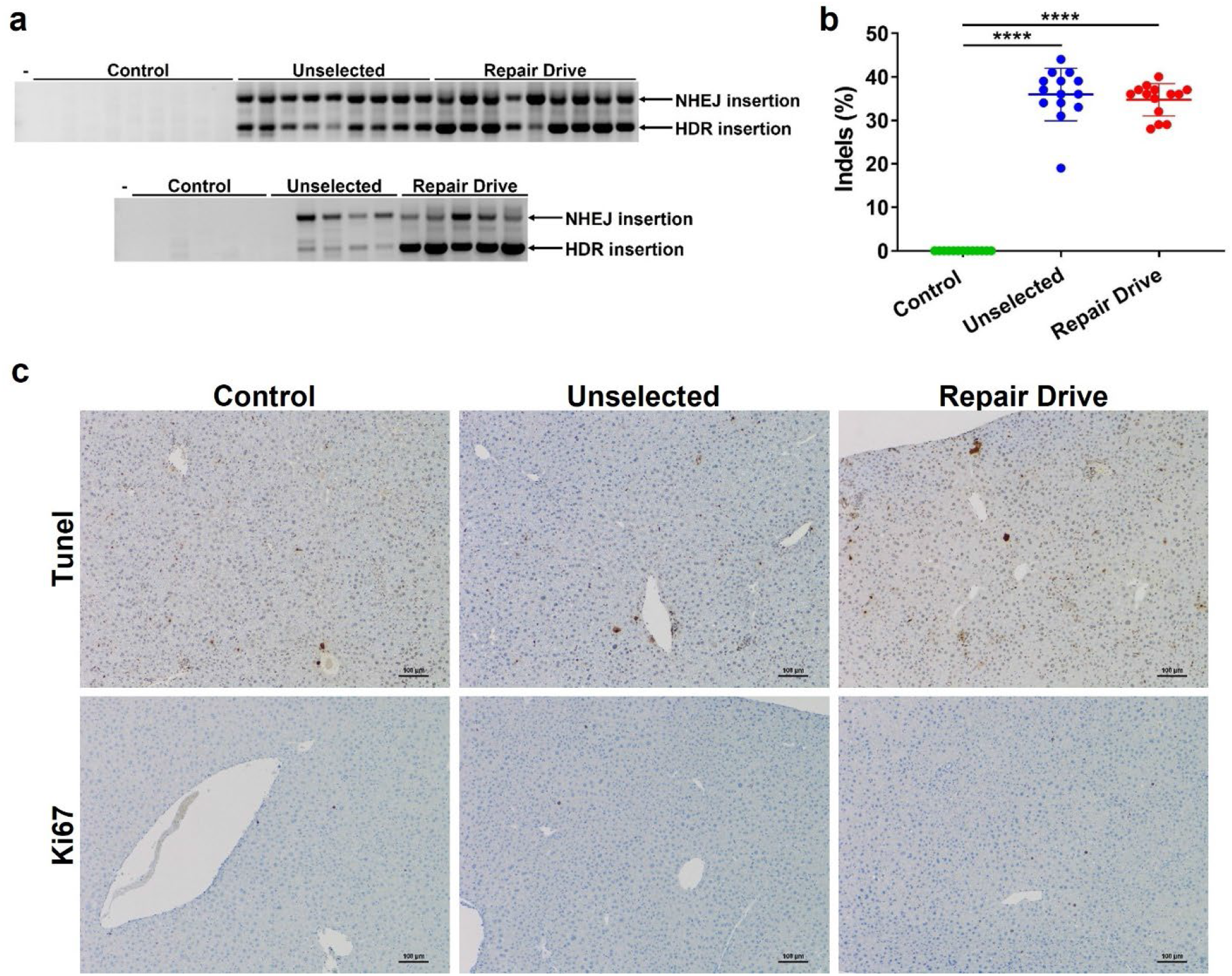
Editing at the *Apoa1* locus and liver histology at one-year post-AAV injection. **a**, PCR from liver show integration of AAV-Donor into the *Apoa1* locus. Two main products were observed: correct HDR (886 bp) and NHEJ insertion (1,778 bp). The forward primer binds to the *Apoa1* locus upstream of the 5’ HR and reverse primer binds to the *FAH* coding sequence. Minus (-) indicates water-only control. **b**, Frequency of indel formation at the *Apoa1* 3’ UTR by ICE analysis. **c**, Representative Tunel and Ki67 staining in livers from Control, Unselected and Repair Drive mice. Scale bar is 100 µm. Data are expressed as mean ± s.d. (n= 13 Control and 14 Unselected and Repair Drive mice), with significance determined by one-way ANOVA followed by Tukey test. **** p<0.0001.

