## Supplemental Figures for "*In vivo* expansion of gene-targeted hepatocytes through transient inhibition of an essential gene"

*Corresponding authors

Correspondence:

**Supplementary Figures and Figure Legends**

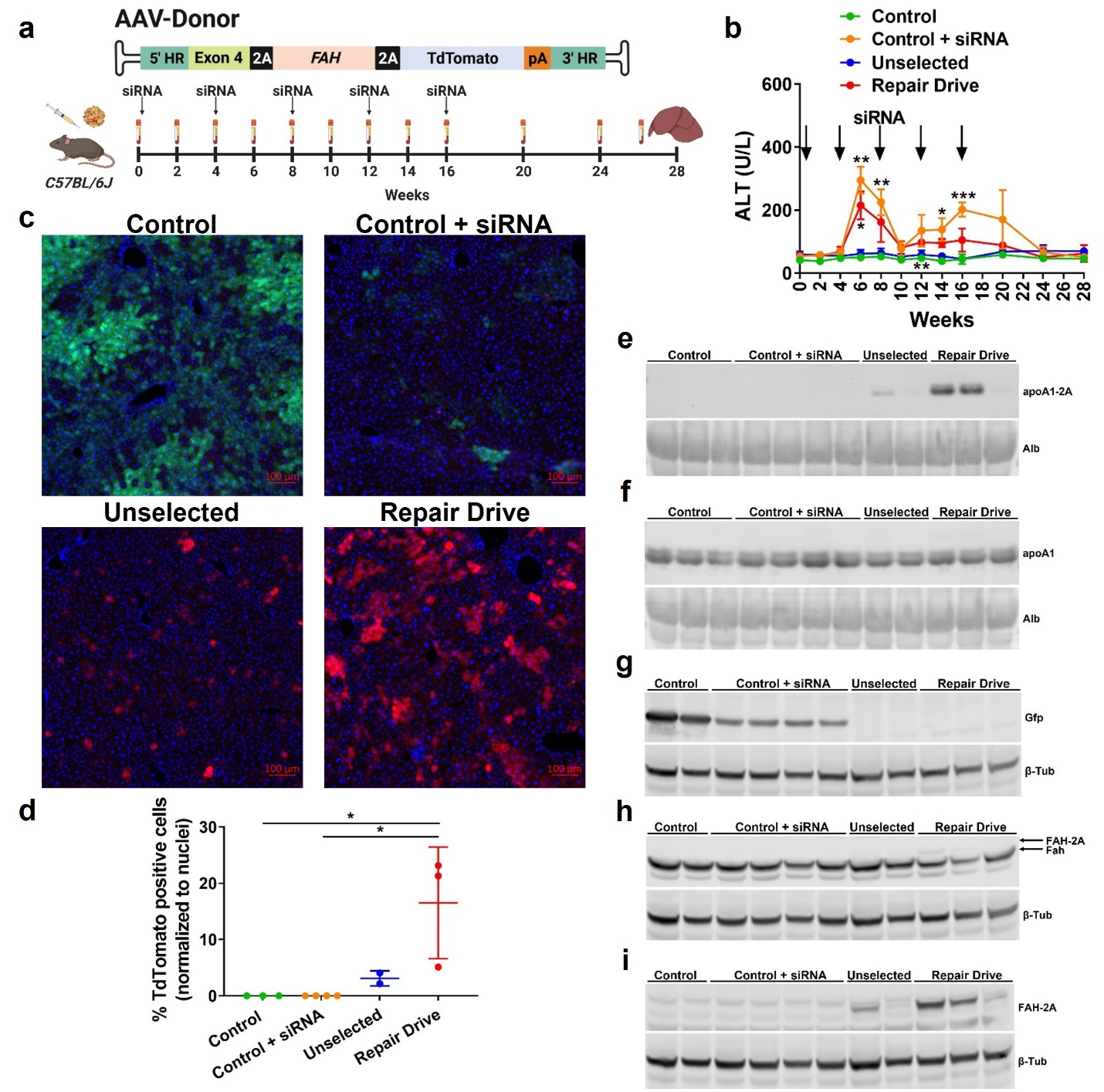

**Supplementary Fig. 1. Pilot test of Repair Drive. a**, Eight-week-old *C57BL/6J* mice were injected with AAVs - 5×10^11^ GC of AAV-CRISPR and 1×10^11^ GC of AAV-Donor or 6×10^11^ GC of AAV-GFP as control. Then, mice were injected with *Fah*-siRNA (3 mg/kg) or saline every four weeks until sixteen weeks post AAV-injection. Mice were sacrificed at 28 weeks post-AAV injection. Blood was collected at time 0 and every two to four weeks. Created with BioRender.com. **b**, Plasma Alt measurement showing transient elevation in Control + siRNA and Repair Drive mice. **c**, Representative direct fluorescence microscopy showing GFP and TdTomato respectively in Control and AAV-CRISPR/Donor-injected mice subjected or no to Repair Drive. Scale bar is 100 µm. **d**, Quantification of TdTomato-positive hepatocytes relative to total nuclei per field. Western blot analysis of 2A-tagged (**e**) and total apoA1 (**f**) in plasma with Alb as loading control. Western blot analysis of Gfp (**g**), endogenous Fah (**h**) and FAH-2A (**i**) in liver lysates with β-Tub as loading control. Data are expressed as mean ± s.d. (n= 2 to 4 mice per group), with significance determined by one-way and two-way ANOVA followed by Tukey test, respectively in **d** and **b**. * p<0.05, ** p<0.01, *** p<0.001 and **** p<0.0001. In (**b**), * p<0.05 Repair Drive vs. Unselected, and ** p<0.01 Control + siRNA vs. Control and Repair Drive at 6 weeks; ** p<0.01 Control + siRNA vs. Control and Unselected at 8 weeks; ** p<0.01 Repair Drive vs. Control at 12 weeks; * p<0.05 Repair Drive vs. Control, and * p<0.05 Control + siRNA vs. Control at 14 weeks; *** p<0.001 Control + siRNA vs. Control and Unselected at 16 weeks.

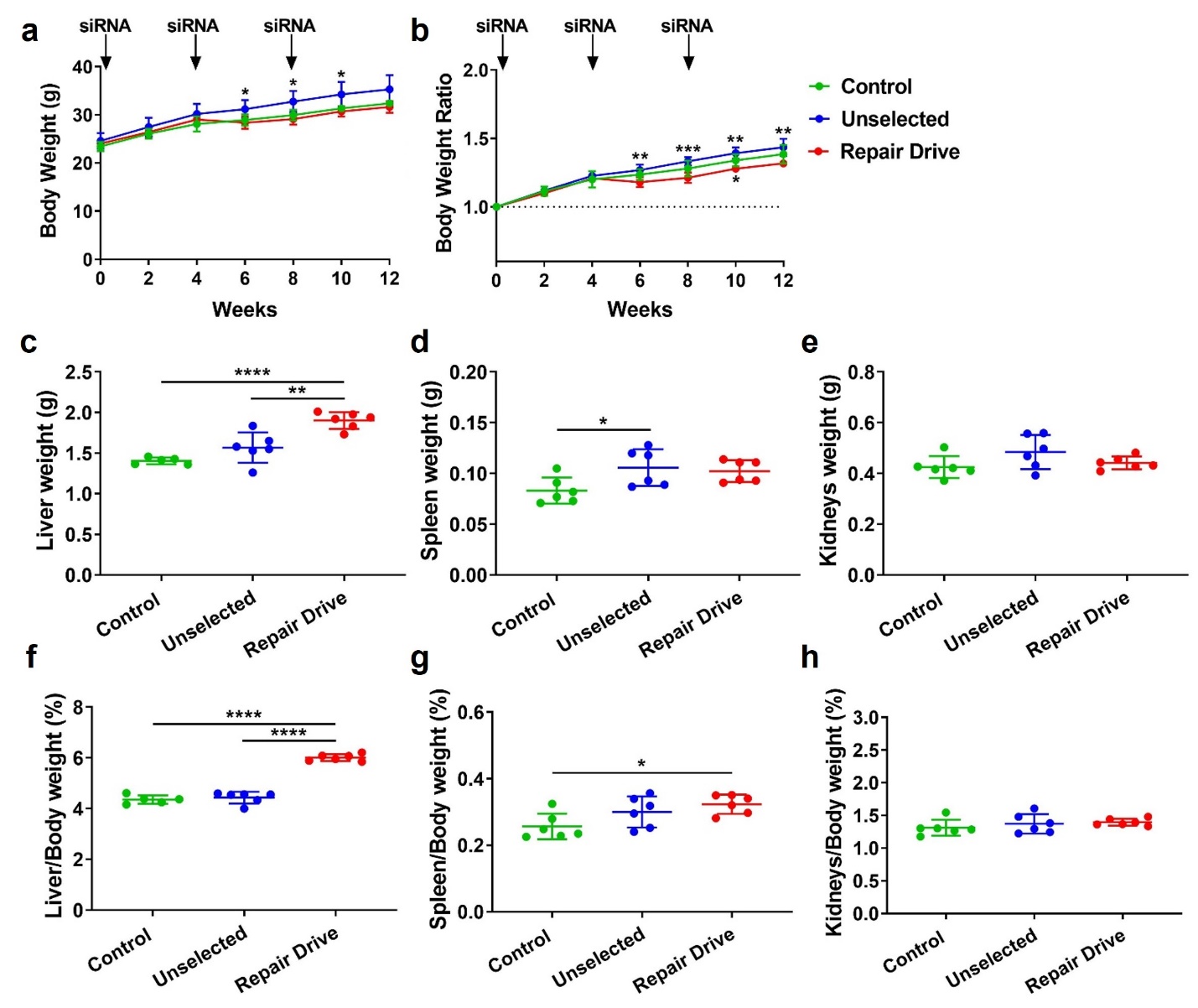

**Supplementary Fig. 2. Body and organ weights in mice subjected to Repair Drive and controls.** Body weights (**a**) and body weight ratios normalized to time 0 (**b**) over time. Endpoint weights of liver (**c**), spleen (**d**) and kidneys (**e**). Endpoint organ to body weight ratios for liver (**f**), spleen (**g**) and kidneys (**h**). Data are expressed as mean ± s.d. (n= 6 mice per group), with significance determined by one-way and two-way ANOVA followed by Tukey test, respectively in **c**-**h** and **a**-**b**. * p<0.05, ** p<0.01, *** p<0.001 and **** p<0.0001. In (**a**), * p<0.05 Repair Drive vs. Unselected at 6, 8 and 10 weeks. In (**b**), ** p<0.01 Repair Drive vs. Unselected at 6 weeks; *** p<0.001 Repair Drive vs. Unselected at 8 weeks; * p<0.05 Repair Drive vs. Control and ** p<0.01 Repair Drive vs. Unselected at 10 weeks; and ** p<0.01 Repair Drive vs. Unselected at 12 weeks.

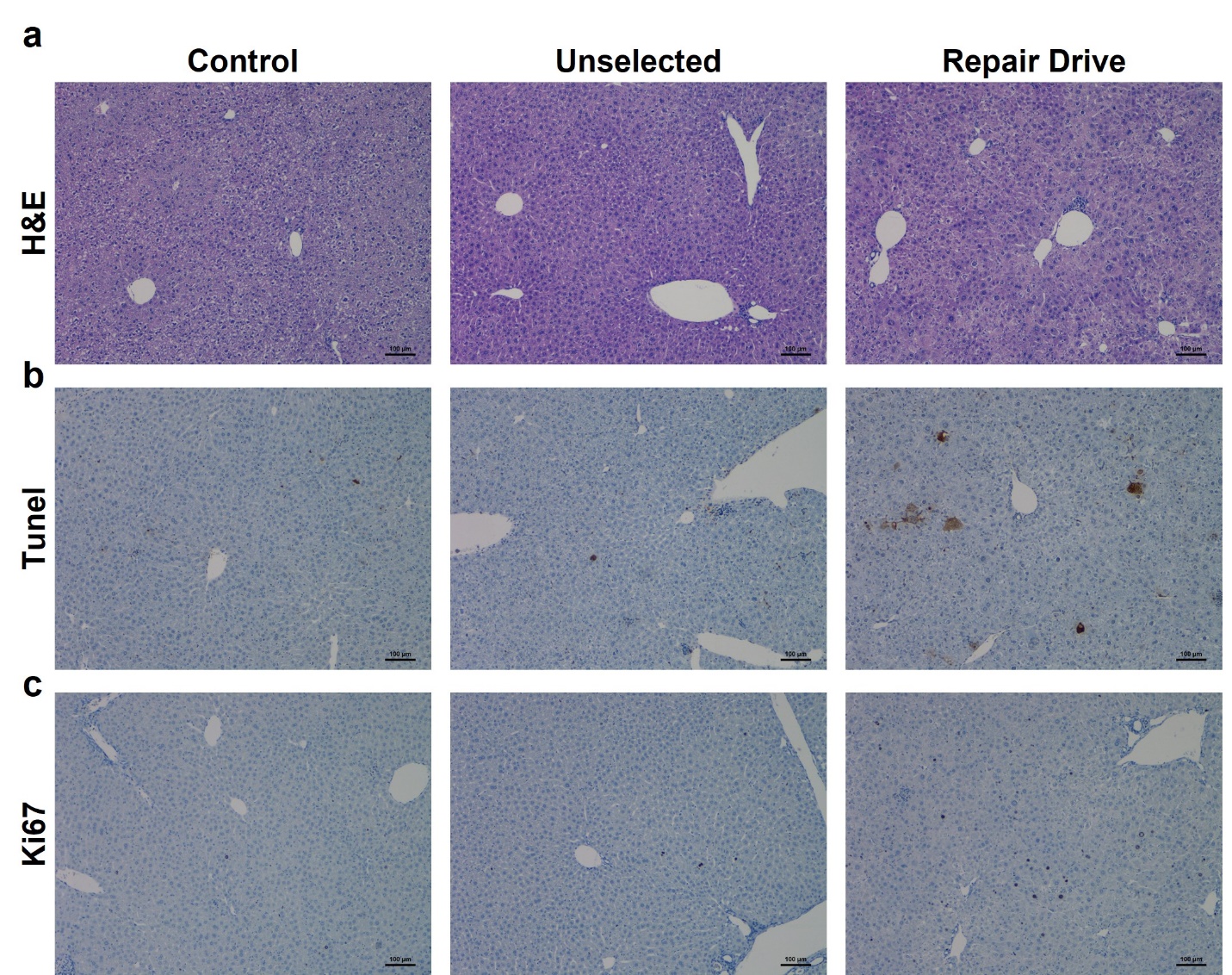

**Supplementary Fig. 3. Hepatocyte cell death and liver regeneration in mice treated with the *Fah*-siRNA.** Representative H&E (**a**), Tunel (**b**) and Ki67 (**c**) staining in livers from Control, Unselected and Repair Drive mice. Scale bar is 100 µm.

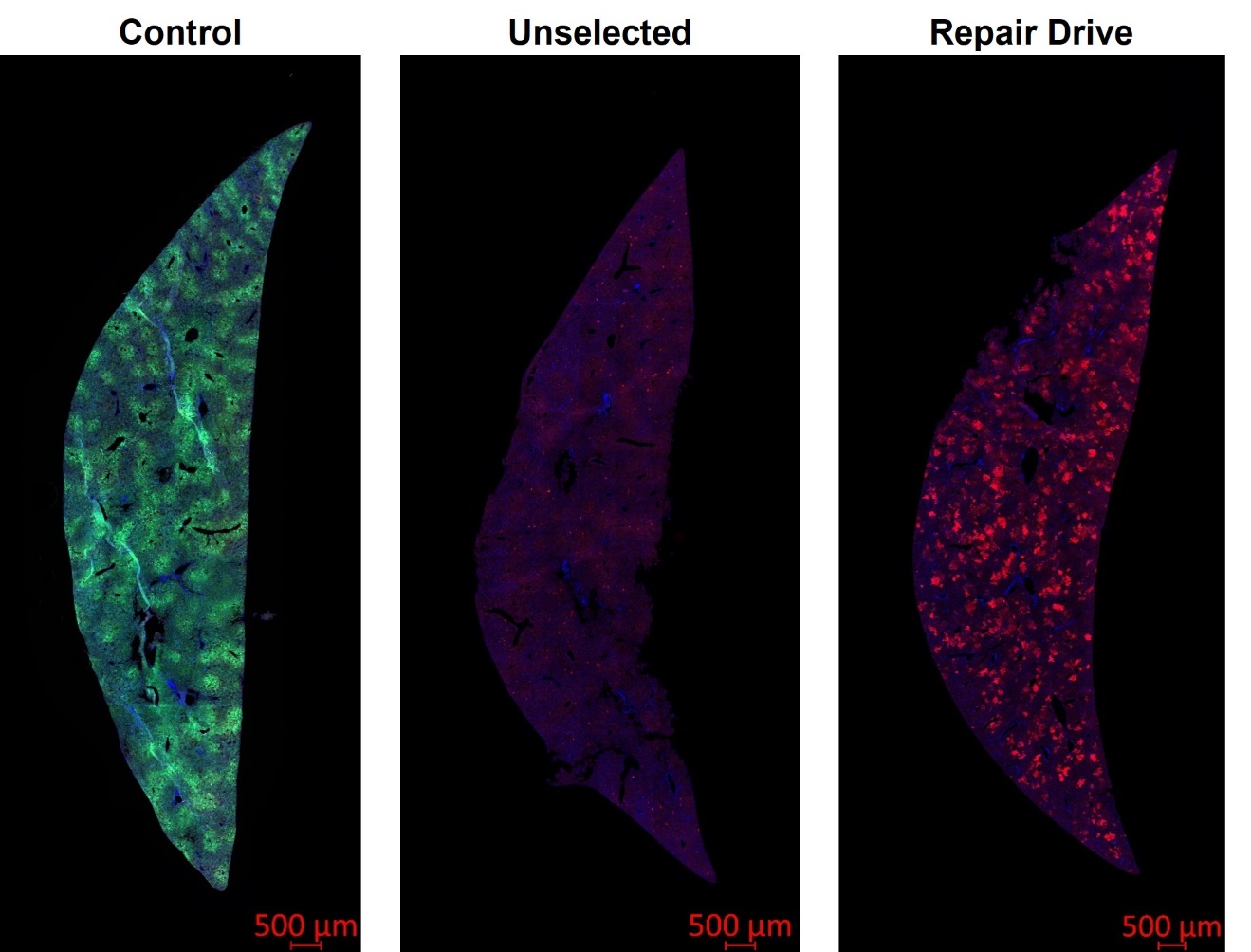

**Supplementary Fig. 4. Enrichment of TdTomato-positive hepatocytes following Repair Drive.** Representative direct fluorescence microscopy of GFP and TdTomato in whole liver sections from Control and AAV-CRISPR/Donor injected mice subjected or no to Repair Drive. Scale bar is 500 µm.

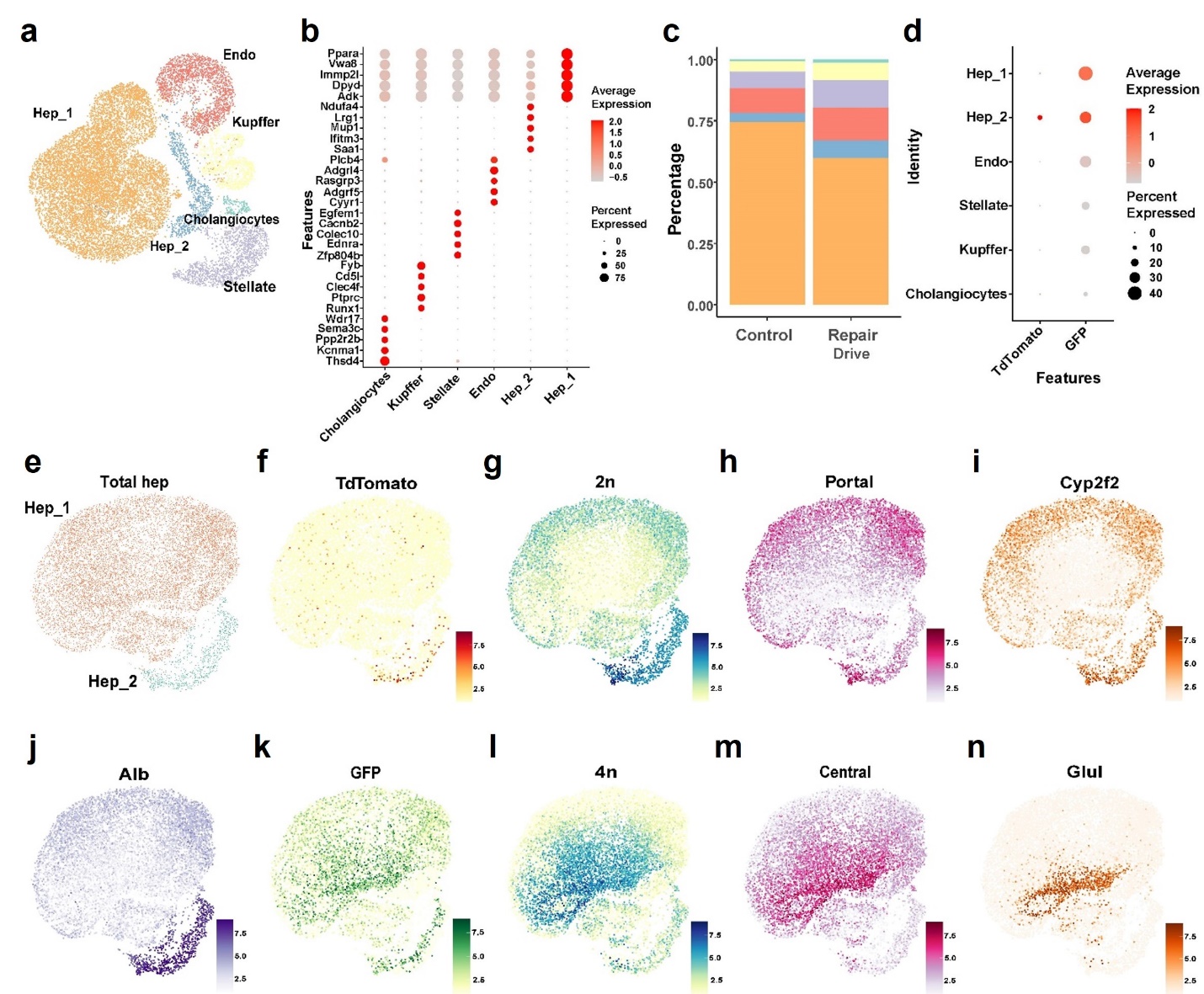

**Supplementary Fig. 5. snRNA-seq analysis of livers from Control and Repair Drive mice. a**, Uniform Manifold Approximation and Projection (UMAP) plot showing the cell types identified in livers from a Control and a Repair Drive mouse: hepatocyte cluster 1 (Hep_1), hepatocyte cluster 2 (Hep_2), endothelial cells (Endo), Kupffer cells, cholangiocytes and stellate cells. **b**, Dotplot showing the expression of marker genes across cell types in the liver. **c**, Cell type abundance in Control vs Repair Drive livers. **d**, Dotplot showing the distribution of TdTomato and GFP expression across cell types in the liver. **e**, UMAP plot showing the two clusters of hepatocytes. UMAP plots showing TdTomato (**f**), diploidy (2n) markers (**g**), portal zonation markers (**h**), Cytochrome P450 2f2 (*Cyp2f2*) (**i**), Alb (**j**), GFP (**k**), tetraploidy (4n) markers (**l**), central zonation markers (**m**) and Glutamate-ammonia ligase (*Glul*) (**n**) expression across Hep_1 and Hep_2 clusters.

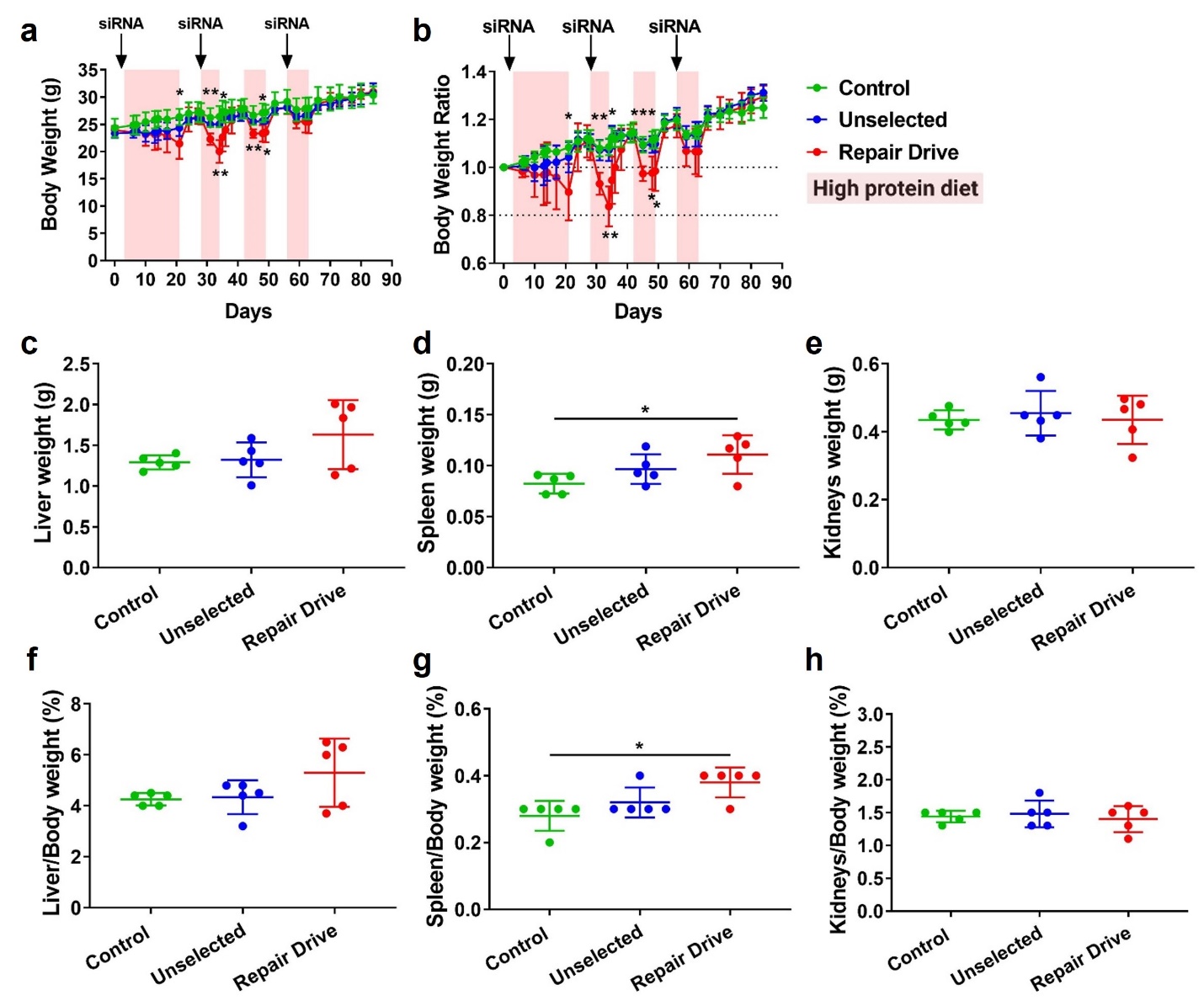

**Supplementary Fig. 6. Body and organ weights in mice subjected to Repair Drive with high protein diet.** Body weights (**a**) and body weight ratios (**b**) normalized to time 0 over time. Endpoint weights of liver (**c**), spleen (**d**) and kidneys (**e**). Endpoint organ to body weight ratios for liver (**f**), spleen (**g**) and kidneys (**h**). Data are expressed as mean ± s.d. (n= 6 mice per group), with significance determined by one-way and two-way ANOVA followed by Tukey test, respectively in **c**-**h** and **a**-**b**. * p<0.05, ** p<0.01 and *** p<0.001. In (**a**), * p<0.05 Repair Drive vs. Control at 21 days; ** p<0.01 Repair Drive vs. Control and Unselected at 31 days; ** p<0.01 Repair Drive vs. Control and Unselected at 34 days; * p<0.05 Repair Drive vs. Control at 35 days; * p<0.05 Repair Drive vs. Control and ** p<0.01 Repair Drive vs. Unselected at 45 days; * p<0.05 Repair Drive vs. Control at 48 days; and * p<0.05 Repair Drive vs. Control at 49 days. In (**b**), * p<0.05 Repair Drive vs. Control at 21 days; ** p<0.01 Repair Drive vs. Control and Unselected at 31 days; ** p<0.01 Repair Drive vs. Control and Unselected at 34 days; * p<0.05 Repair Drive vs. Control and Unselected at 35 days; *** p<0.001 Repair Drive vs. Control and Unselected at 45 days; * p<0.05 Repair Drive vs. Control and Unselected at 48 days; and * p<0.05 Repair Drive vs. Control and Unselected at 49 days.

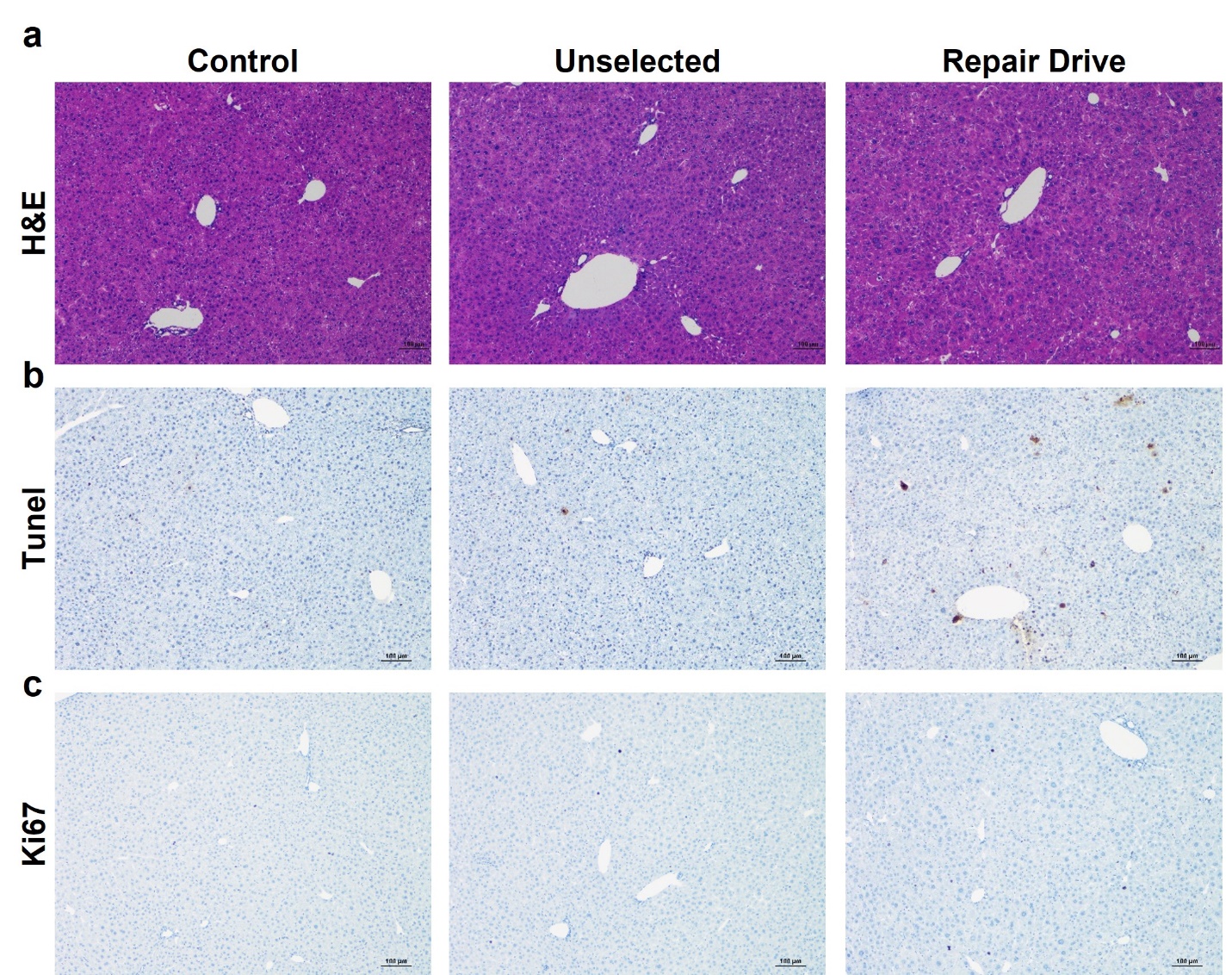

**Supplementary Fig. 7. Hepatocyte cell death and liver regeneration in mice treated with the *Fah*-siRNA and fed a high protein diet.** Representative H&E (**a**), Tunel (**b**) and Ki67 (**c**) staining in livers from Control, Unselected and Repair Drive mice. Scale bar is 100 µm.

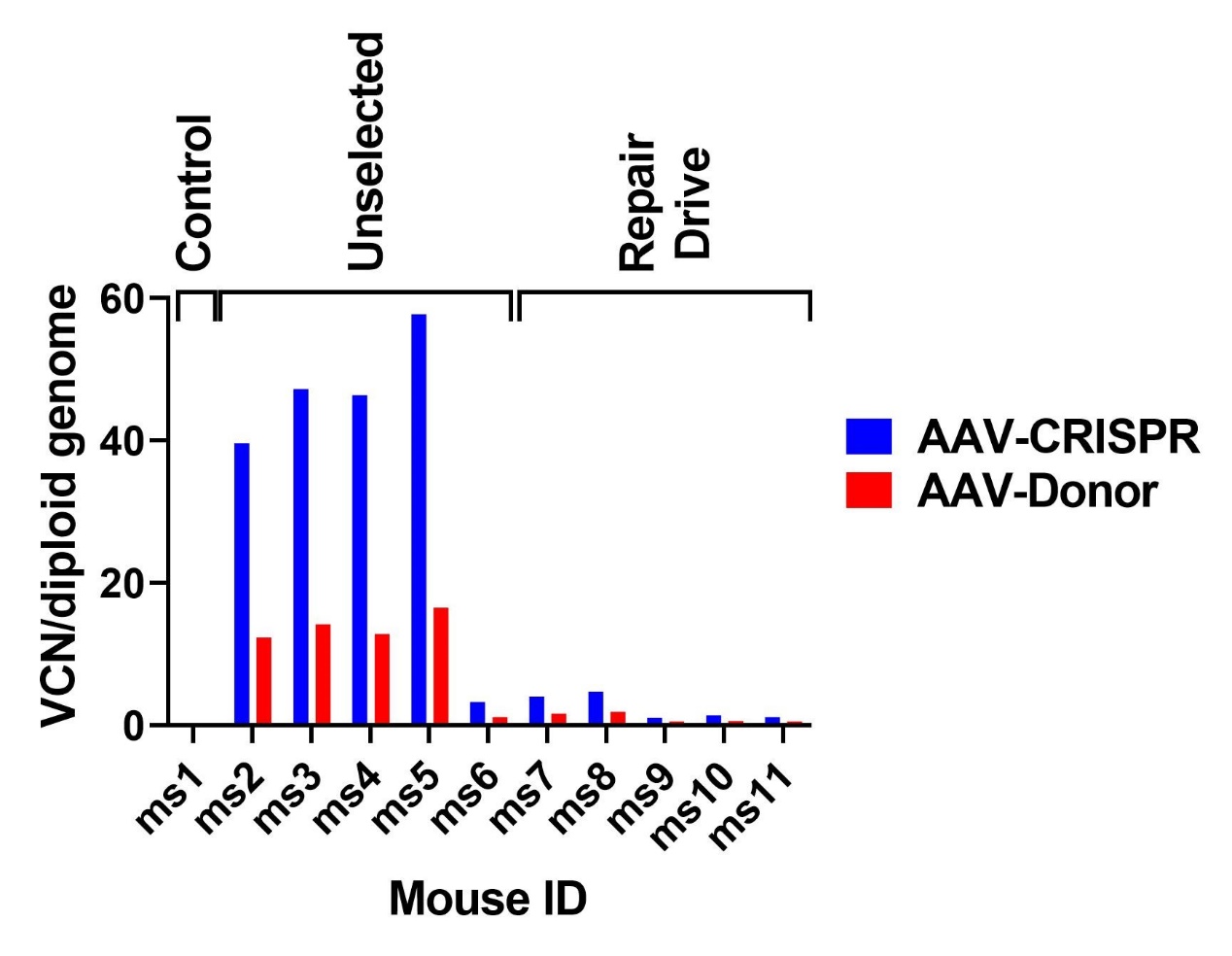

**Supplementary Fig. 8. Depletion of episomal AAVs following Repair Drive.** Vector copy number (VCN) per diploid genome of AAV-CRISPR and AAV-Donor measured by ddPCR in Unselected and Repair Drive livers. Ms: mouse.

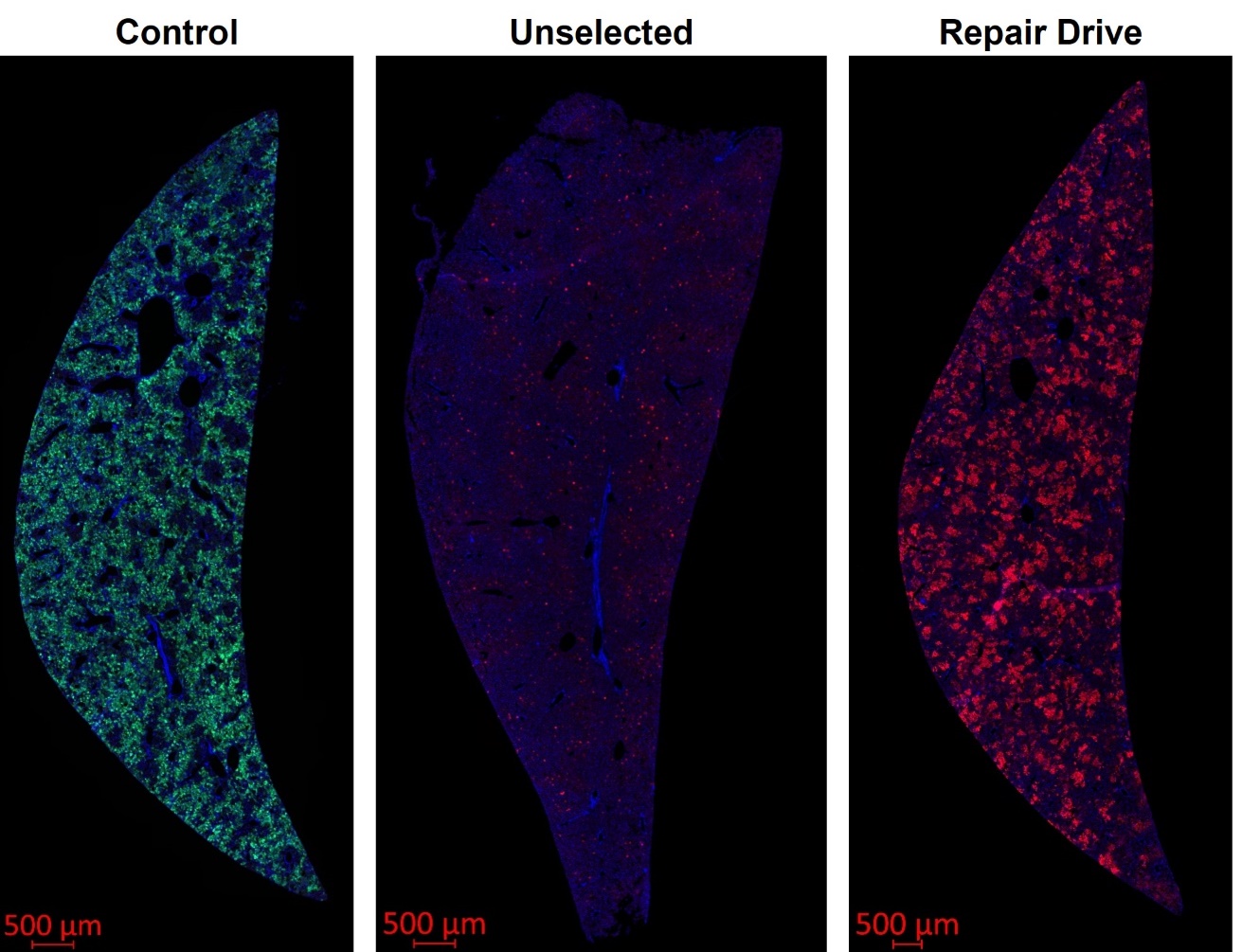

**Supplementary Fig. 9. Enrichment of TdTomato-positive hepatocytes following Repair Drive with high protein diet.** Representative direct immunofluorescence microscopy of GFP and TdTomato in whole liver sections from Control and AAV-CRISPR/Donor injected mice subjected or no to Repair Drive. Scale bar is 500 µm.

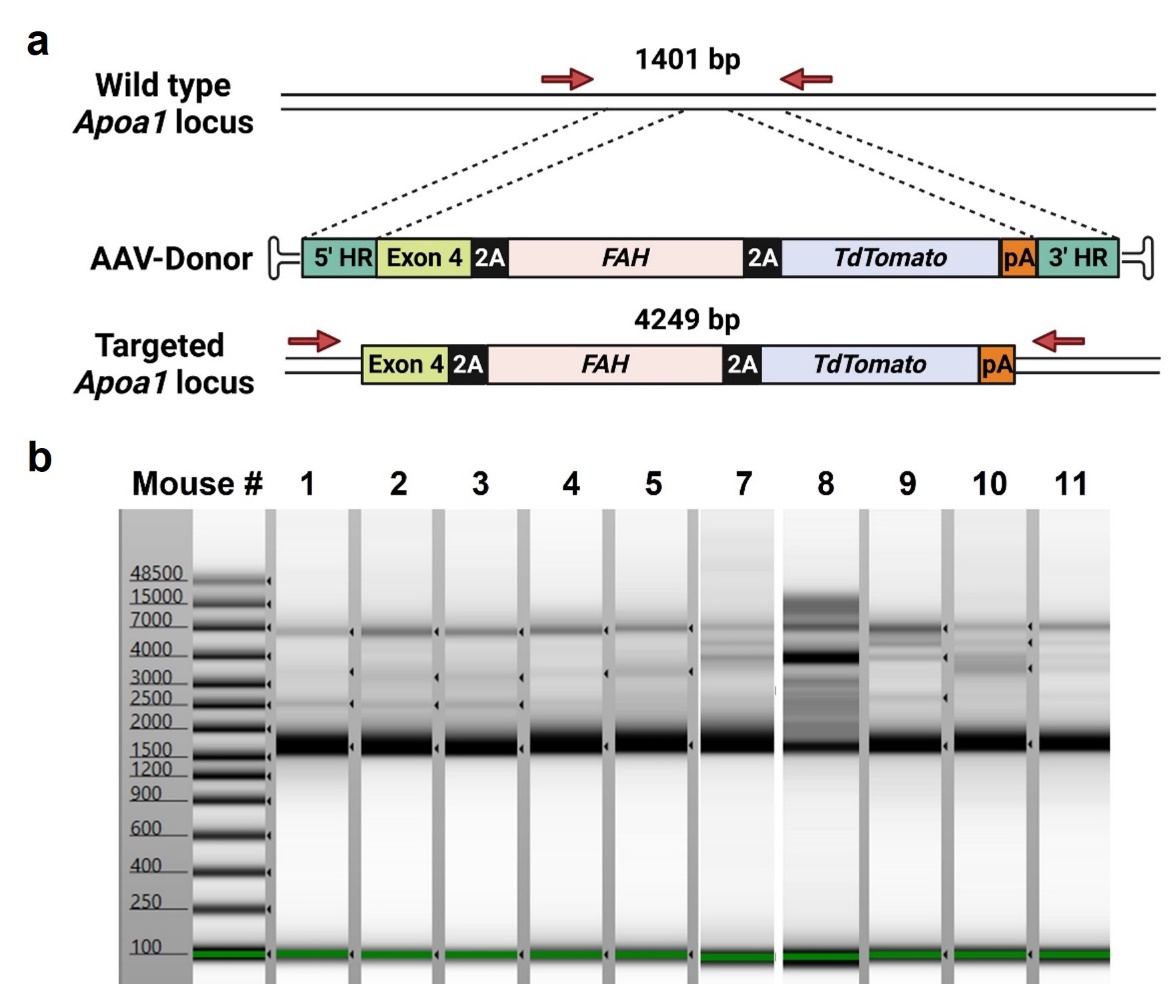

**Supplementary Fig. 10. Sample preparation for SMRT-seq with UMI. a**, Scheme of long-range PCR of the *Apoa1* locus with expected amplicons generated starting from unmodified (1401 bp) and HDR alleles (4249 bp). Created with BioRender.com. **b**, DNA bioanalyzer image of dual-UMI labeled long-range PCR products. Mouse 1: Control; Mouse 2-5: Unselected mice; Mouse 7-11: Repair Drive mice. The ~ 1500 bp PCR product derives from the amplification of wild type *Apoa1* alleles or *Apoa1* alleles with small indels. The low intensity higher molecular weight PCR products (up to 7000 bp) derive from *Apoa1* alleles with different outcomes of AAV integration. Mouse 6 was sequenced using SMRT-seq with UMI, but not included in the gel. In mouse 1, the faint high molecular weight PCR products are likely artifacts as shown by the absence of UMI consensus reads with insertions.

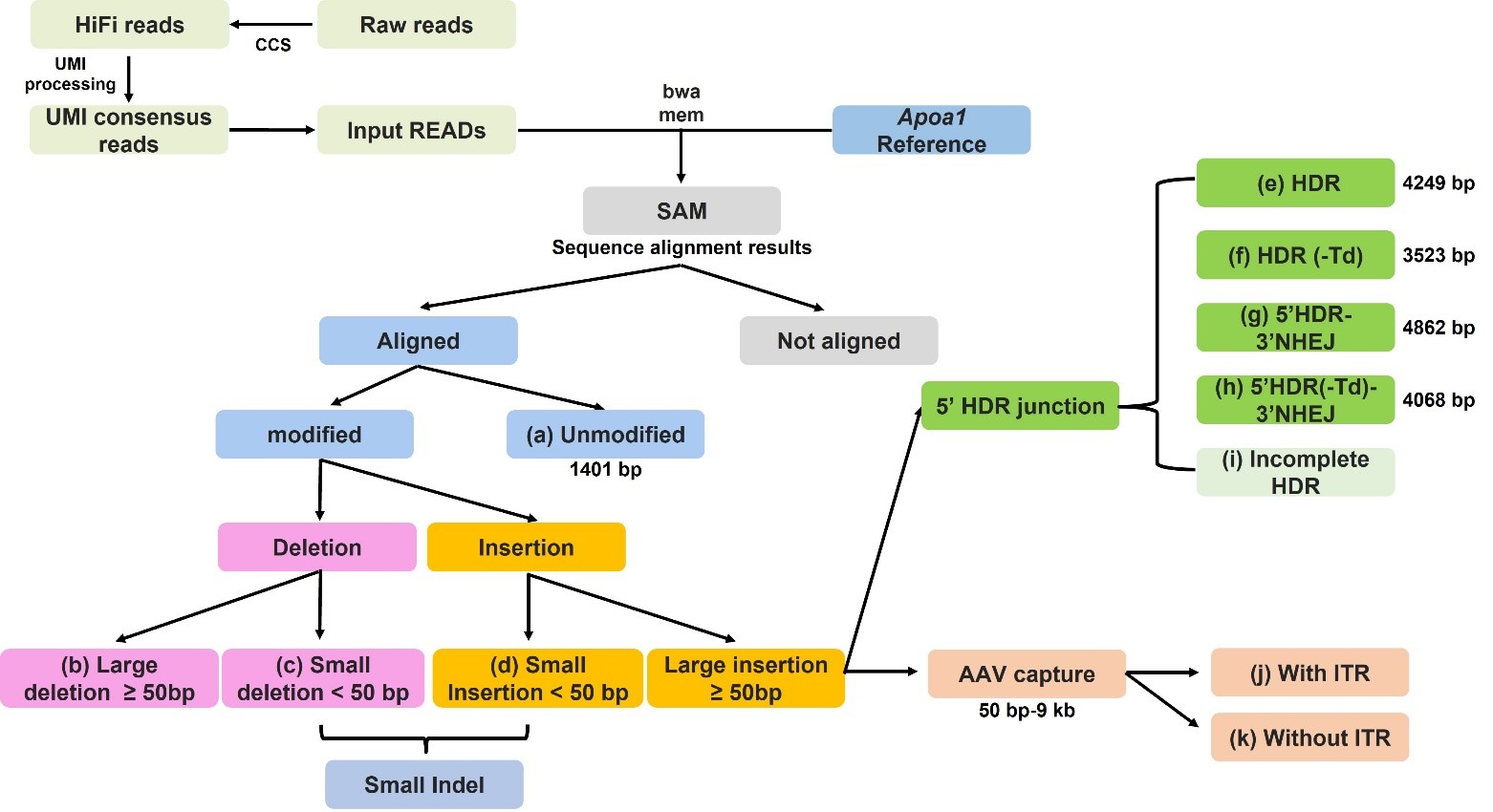

**Supplementary Fig. 11. Schematics of SMRT-seq with UMI processing and variant calling pipeline.** Each PCR products of 1-6 kb around the Cas9 cut site was tagged with dual UMIs. The longread_umi pipeline described by Karst et al. was adapted to generate UMI consensus sequences from demultiplexed PacBio CCS reads.^1^ UMI consensus sequences were aligned to the reference amplicon sequence using Burrows-Wheeler Alignment (BWA)-Maximal Exact Match (MEM).^2^ We used a custom Matlab script to analyze the UMI consensus sequences and characterize the gene editing outcomes. In short, gene modification variants were categorized into ten groups: (a) unmodified (1401 bp), (b) large deletions (≥ 50 bp), (c) small deletion (< 50 bp), (d) small insertion (< 50 bp), (e) HDR (4249 bp), (f) HDR with truncation of one tdTomato – HDR(-Td) (3523 bp), (g) 5’HDR-3’NHEJ (4862 bp), (h) 5’HDR(-Td)-3’NHEJ (4068 bp), (i) incomplete HDR with partial integration of insert, (j) AAV capture (50 bp – 9 kb) with ITR at the integration junction and (k) ITR-less AAV capture.

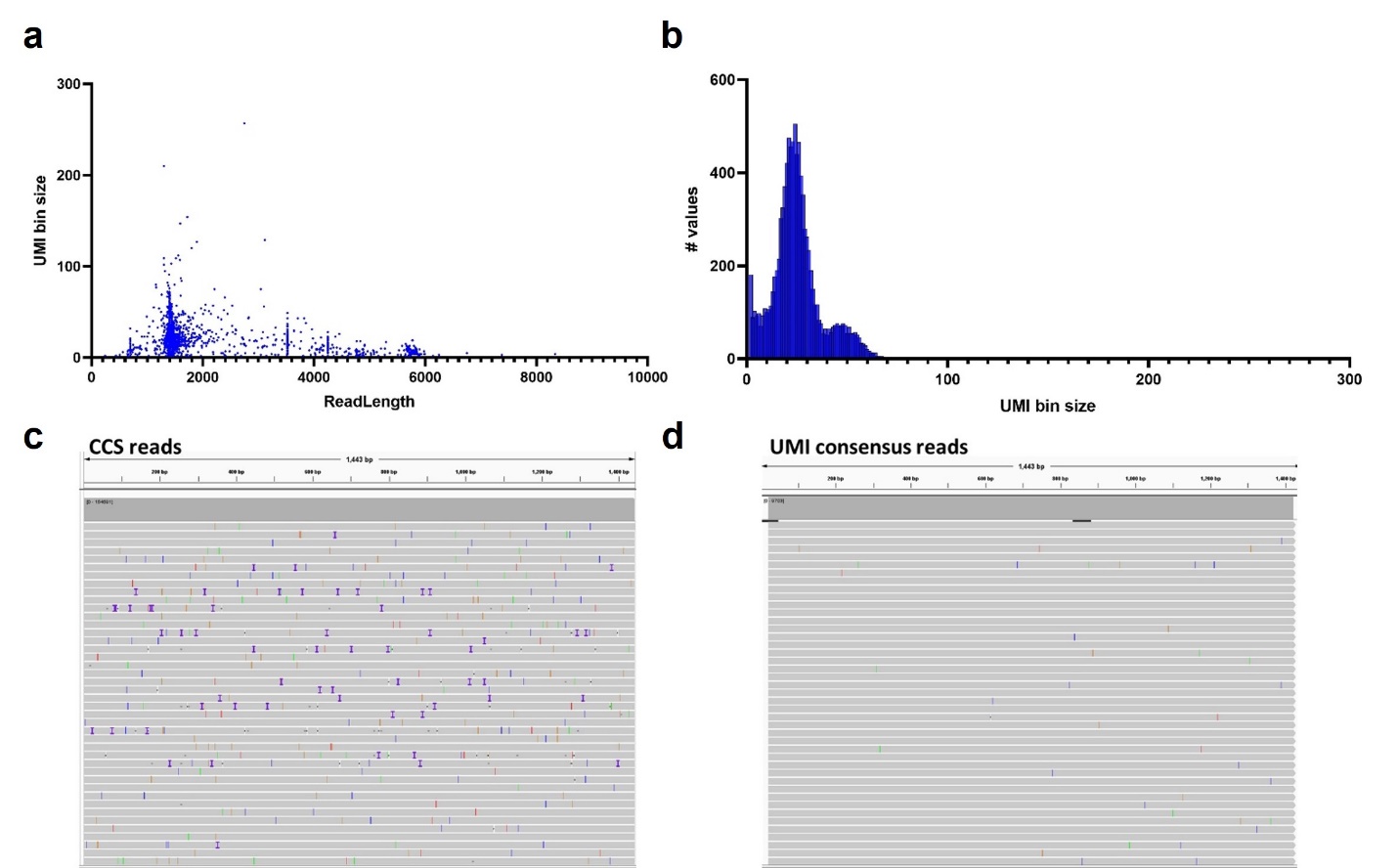

**Supplementary Fig. 12. High accuracy UMI consensus reads. a**, UMI cluster size for individual CCS reads from mouse 11 (Repair Drive). **b**, UMI cluster size histogram of sequenced reads from mouse 11. Consensus sequence for each UMI bin (clustered UMI pair) was generated by multiple rounds of polishing using the binned raw reads. UMI pairs with three or more CCS reads were used for UMI consensus sequence generation to remove PCR errors based on the sequence information within each CCS read derived from the same SMRTbell template molecule. For mouse 11, we generated 9783 UMI consensus sequences from 233323 CCS reads. **c**, CCS reads and (**d**) UMI consensus reads from mouse 1 (Control) aligned to *Apoa1* reference sequence and visualized on Integrative Genomics Viewer (IGV), showing highly polished UMI consensus reads after UMI processing and consensus read generation. Vertical colored lines 1 bp indels or mismatches from PCR or sequencing error found within the amplicon compared to the reference sequence.

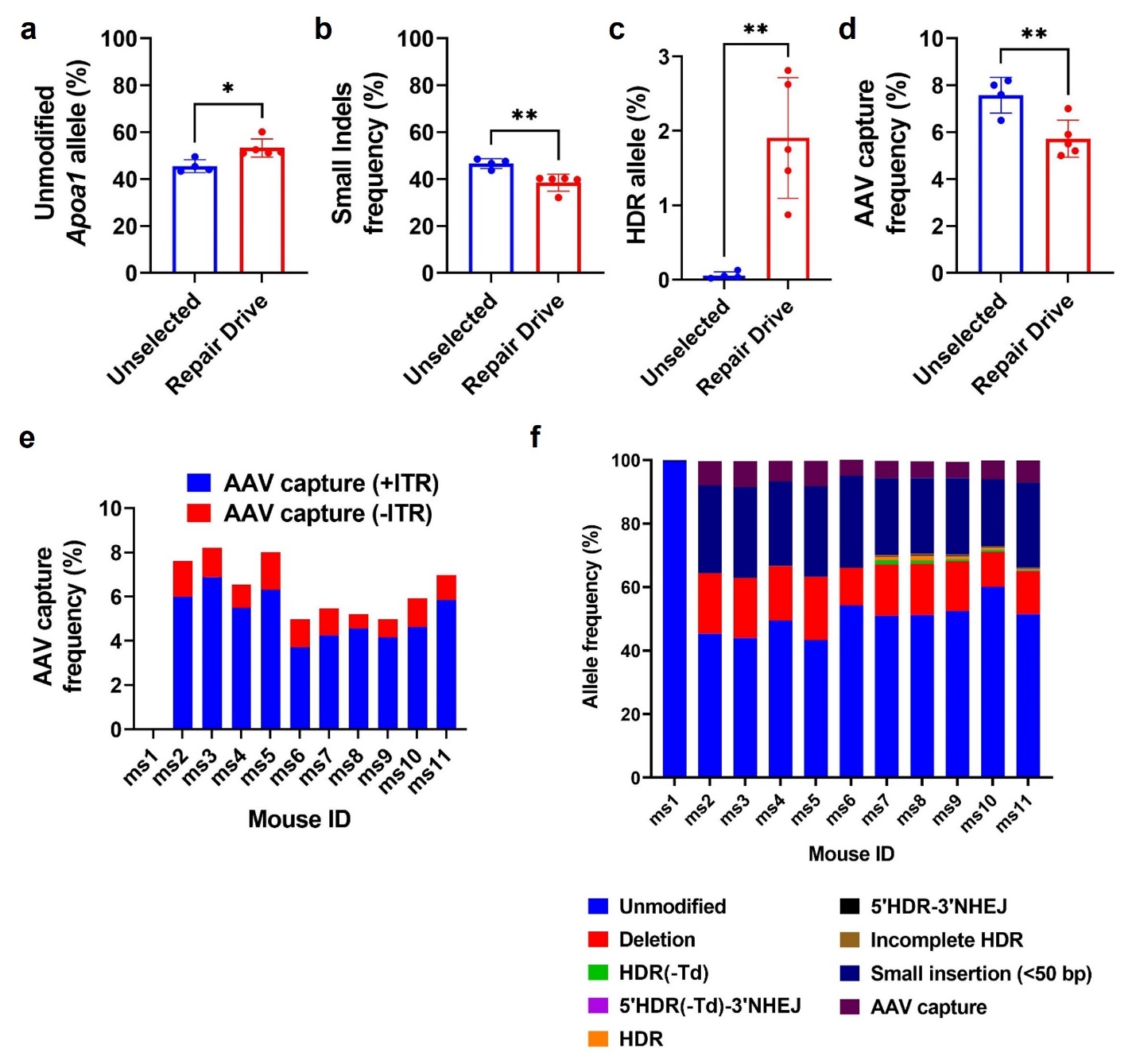

**Supplementary Fig. 13. Editing analysis at the *Apoa1* locus by SMRT-seq. a**, Frequency of unmodified *Apoa1* alleles in Unselected and Repair Drive mice. Frequency of *Apoa1* alleles with (**b**) small indels, (**c**) HDR-integration of AAV-Donor and (**d**) NHEJ-insertion of AAVs in Unselected and Repair Drive mice. **e**, Frequency of NHEJ-insertion of AAVs with or without ITR sequences in Control (ms1), Unselected (ms2 - ms6) and Repair Drive (ms7 – ms11) mice. **f**, Overall frequency of unmodified and differently edited *Apoa1* alleles in Control (ms1), Unselected (ms2 - ms6) and Repair Drive (ms7 – ms11) mice. Data are expressed as mean ± s.d. (n= 1 Control, 5 Unselected and Repair Drive mice) with significance determined by two-tailed student’s t-test. * p<0.05 and ** p<0.01.

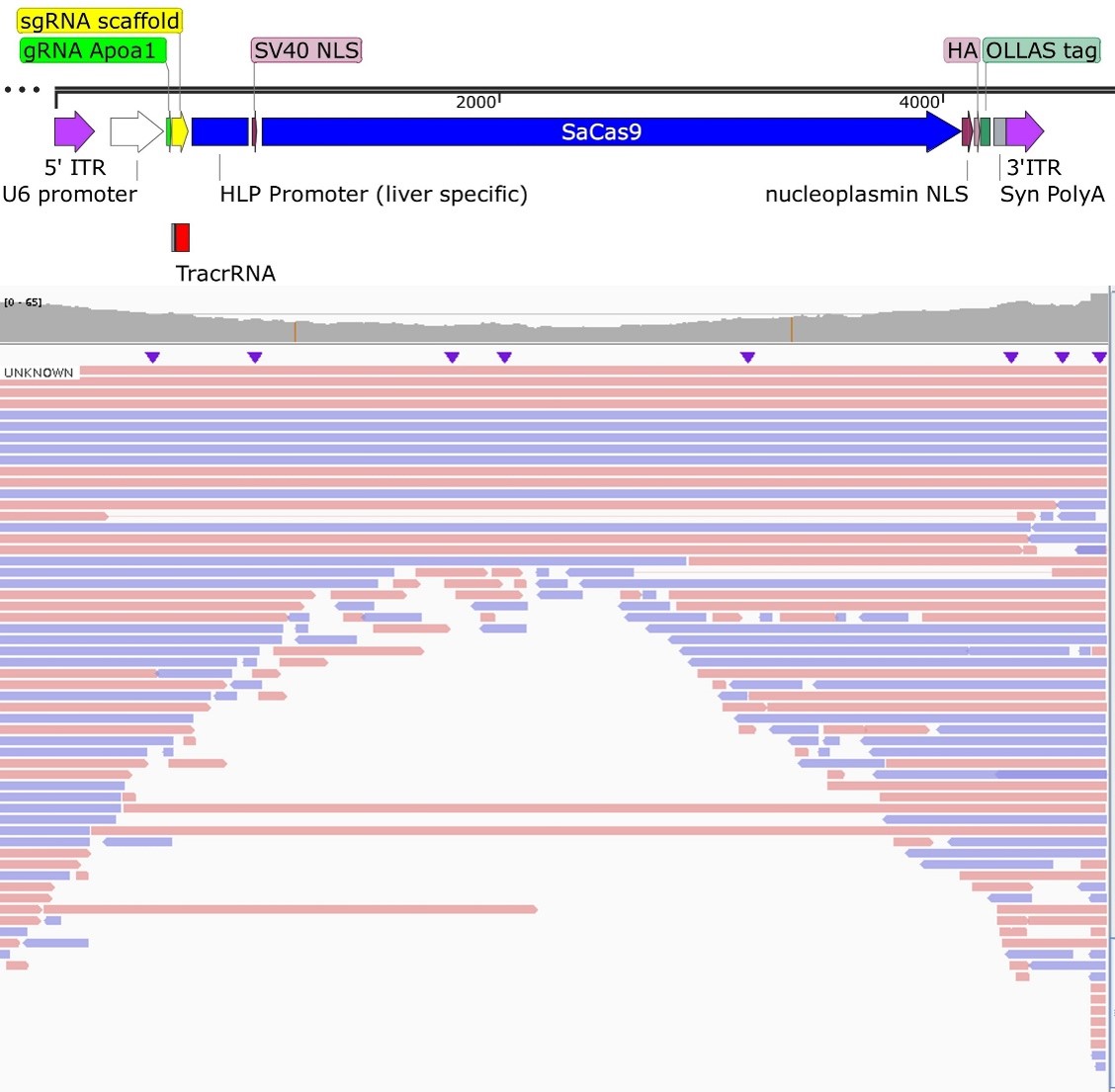

**Supplementary Fig. 14. IGV alignment of on-target AAV capture sequences aligned to AAV-CRISPR.** Most on-target captured AAV sequences were 50 bp – 5 kb fragments showing AAV breakage and subsequent integration from internal vector fragments. We also found full AAV genome capture or AAV concatemer with combinations of head-to-tail or tail-to-tail conformations of AAV-CRISPR and AAV-DONOR. We also found the integration of AAV-CRISPR with sequences from the pEMBL8 AAV plasmid, although at very low abundance.

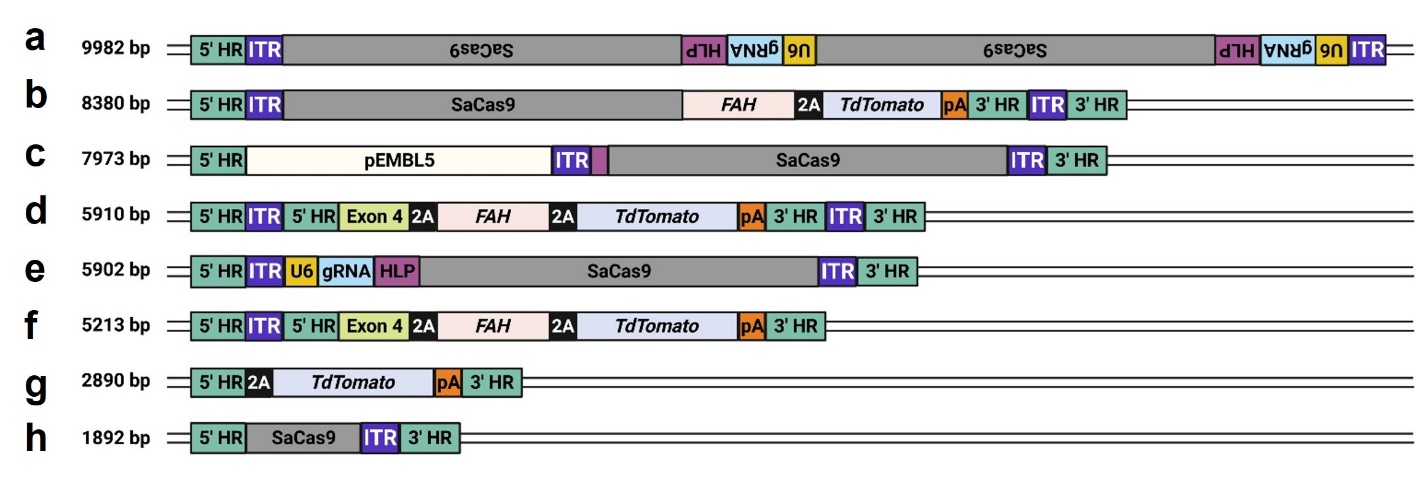

**Supplementary Fig. 15. Schematics of on-target AAV-Donor and AAV-CRISPR capture identified by SMRT-seq with UMI.** Example schematics of AAV capture at the *Apoa1* on-target site including (**a**) Inverted AAV-CRISPR head-to-tail concatemer (9982 bp); (**b**) AAV-CRISPR and AAV-Donor concatemer (8380 bp); (**c**) pEMBL5 AAV packaging plasmid and AAV-CRISPR concatemer (7973 bp); (**d**) NHEJ integration of AAV-Donor (5910 bp); (**e**) NHEJ integration of AAV-CRISPR (5902 bp); (**f**) 5’NHEJ-3’HDR integration of AAV-Donor (5213 bp); (**g**) Partial integration of AAV-Donor (2890 bp); and (**h**) Partial integration of AAV-CRISPR (1892 bp). Created with BioRender.com.

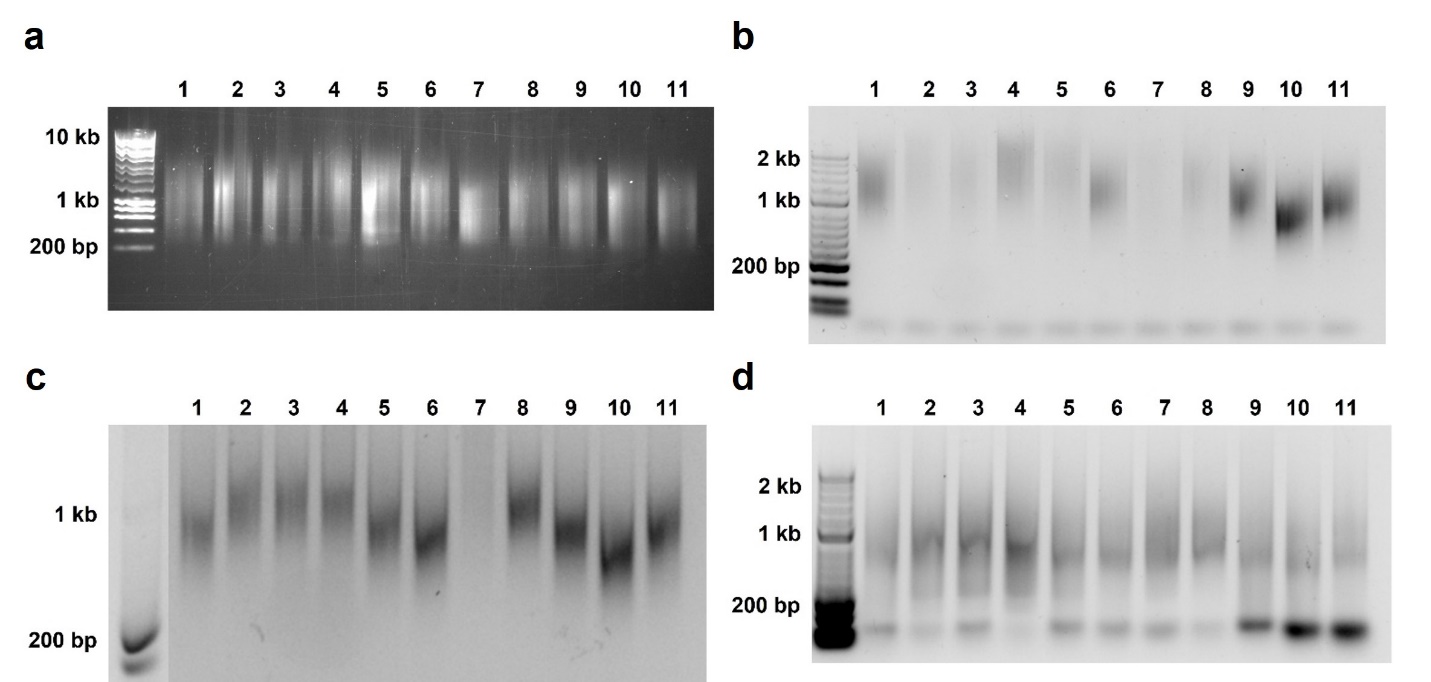

**Supplementary Fig. 16. Sample preparation for GISA-seq.** **a**, DNA was extracted from the liver of eleven mice, including one Control (lane 1), five Unselected (lanes 2-6) and five Repair Drive (lanes 7-11) mice. 4 µg of genomic DNA were randomly fragmented to 300 bp – 6 kb long fragments and visualized on agarose gel. **b**, PCR2 products. After end-repair and A-tailing, unique Y-adapters containing UMI and sample-specific P5 index were ligated to the DNA fragments. Y-adapter-ligated DNA fragments were amplified by PCR1 using a TdTomato-specific forward primer and a reverse primer binding to the P5 sequence on Y-adapter. Subsequent nested PCR2 was performed to enrich the target amplicons and improve specificity. **c**, PCR3 products. PCR3 was performed using staggered forward primer annealing at 3’HR and sample-specific P5 index reverse primer to add adapter sequences and reduce amplicon size. **d**, PCR4 products. PCR4 was performed using index primers to add sample-specific P7 index.

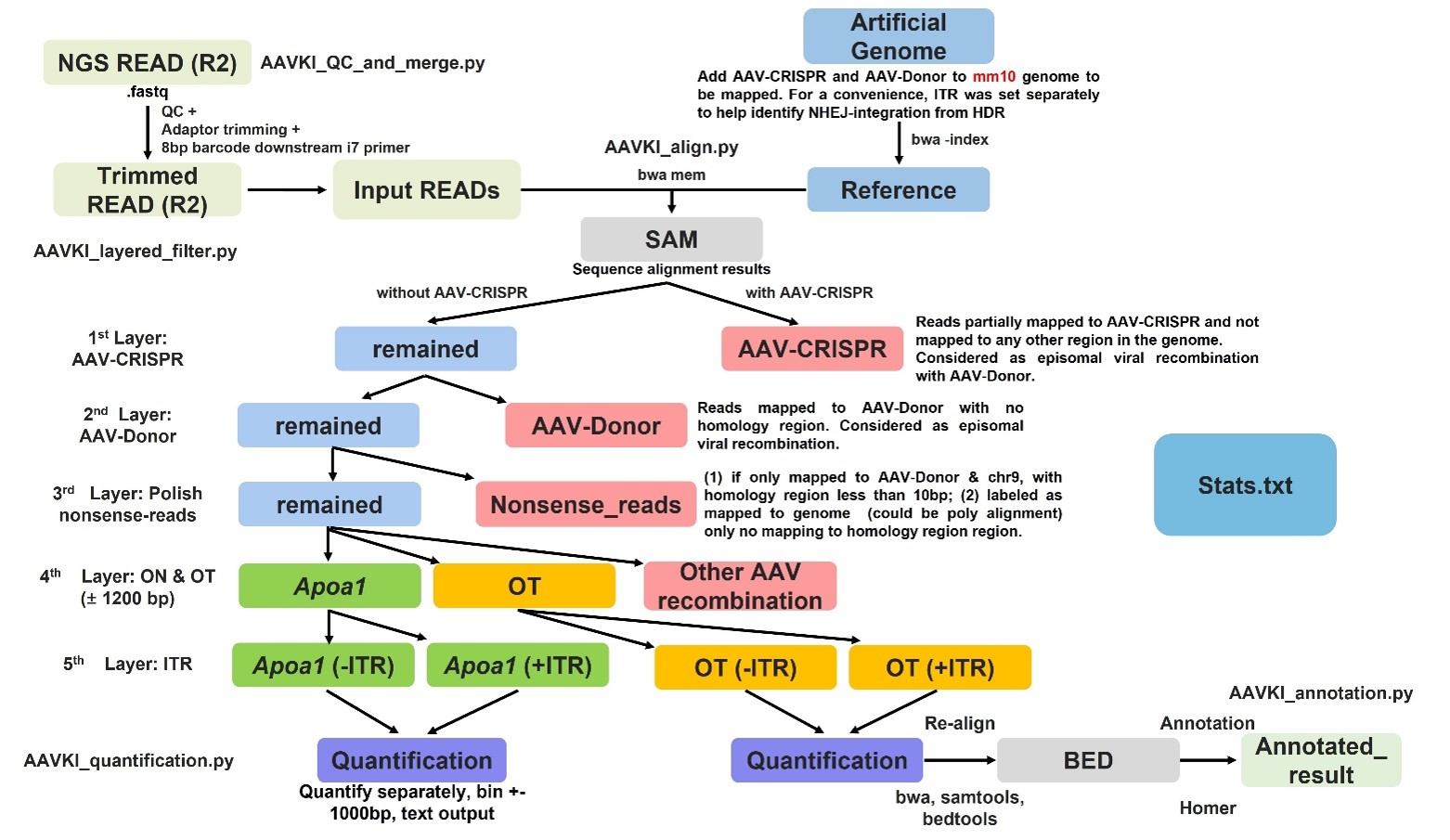

**Supplementary Fig. 17. Scheme of AAV-Donor integration sites identification and quantification assay bioinformatics workflow.** The demultiplexed and consolidated Read2 was processed using an ad-hoc bioinformatics pipeline. A custom reference genome was built, including mm10, ITR-less AAV-CRISPR, ITR-less AAV-Donor, and ITR sequences. Reads generated from non-specific amplification were filtered using a target-specific 8 bp sequence. NGS sequences were aligned against the custom reference genome using BWA-MEM, and reads unable to map properly to the reference were removed during the filtering step. The resulting Read2 contained 53 bp 3’HR followed by the sequence of interest. Multiple filters were applied to remove episomal AAV sequences: (1) AAV-CRISPR filter removed reads mapping to AAV-CRISPR and not to mm10; (2) AAV-Donor filter removed reads mapping to AAV-Donor excluding the 53 bp 3’HR sequence; (3) Complex recombination of AAV-Donor and AAV-CRISPR were detected and removed; the resulting reads had AAV-Donor integration in mm10 and were further categorized into (4) on-target (ON) AAV-Donor integration at the *Apoa1* locus or (5) off-target (OT) AAV-Donor integration. OT AAV-Donor integration reads were further categorized based on the presence or absence of ITR at the integration junction. For ONs, start mapping positions were consolidated using a 100-bp sliding window. For OTs, start mapping positions were consolidated using a 1000-bp sliding window. The sequence was retrieved from each OT integration site and re-aligned to the mm10 using a BWA-MEM to generate BED (Browser Extensible Data) files using Samtools and Bedtools for further annotation using HOMER (Hypergeometric Optimization of Motif EnRichment) suite.

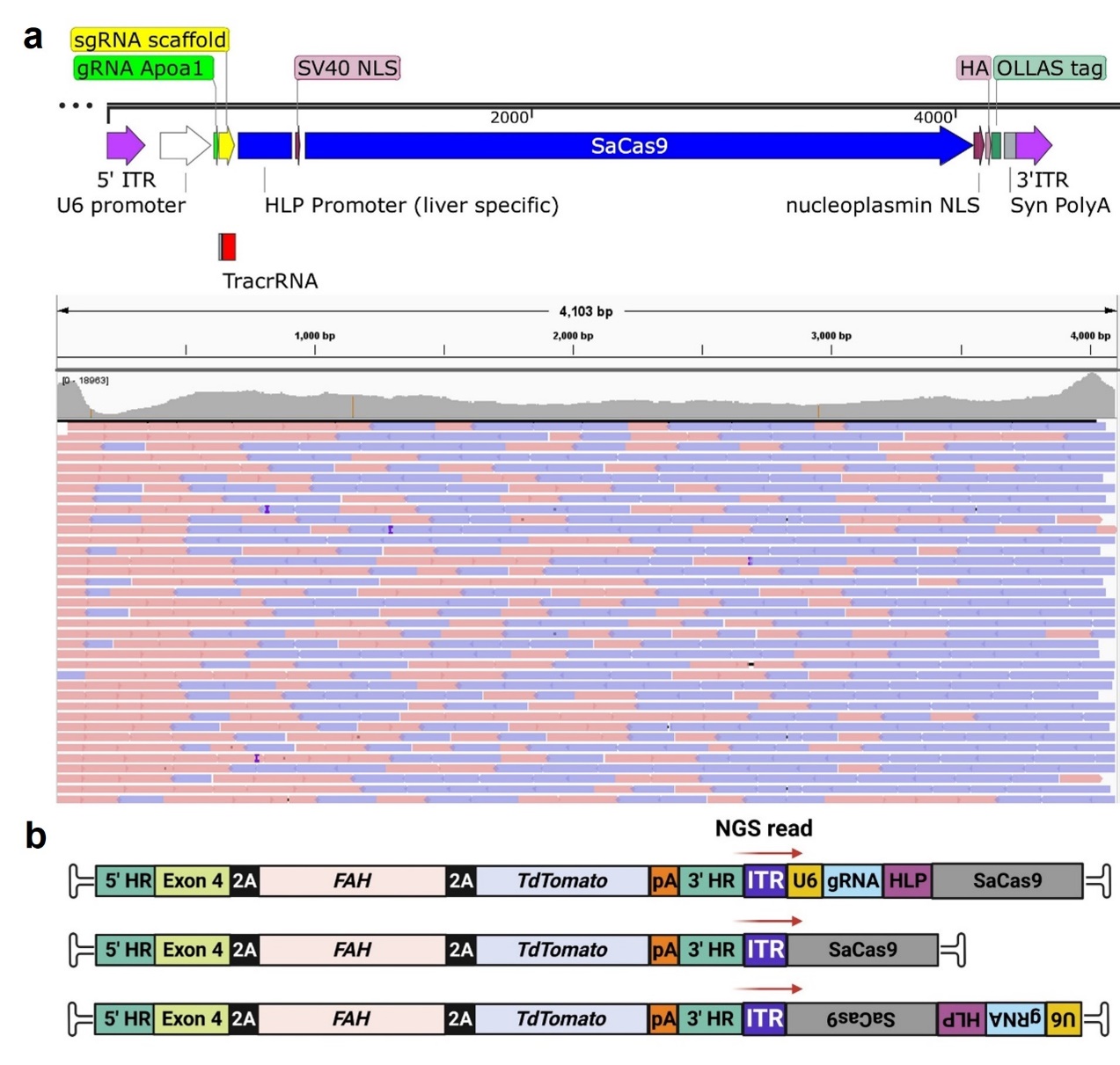

**Supplementary Fig. 18. IVG alignment of GISA-seq reads showing recombination between AAV-Donor and AAV-CRISPR.** **a**, IVG alignments of GISA-seq reads mapping to AAV-CRISPR. These reads were a result of direct ligation of AAV-Donor and AAV-CRISPR in tail-to-head or tail-to-tail conformation as shown in panel **b**. **b** Created with BioRender.com.

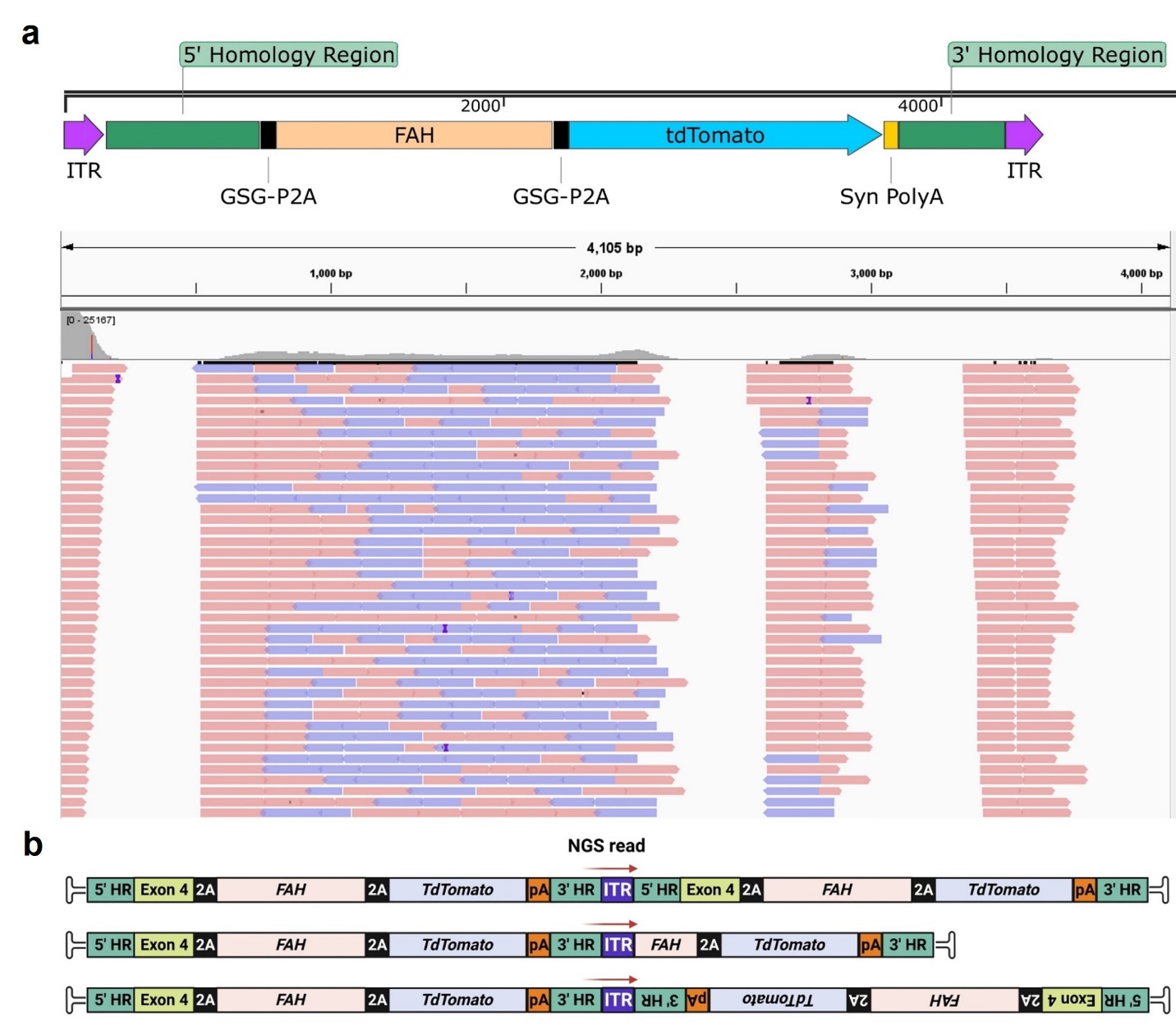

**Supplementary Fig. 19. IVG alignment of GISA-seq reads showing recombination of AAV-Donor.** **a**, IVG alignments of GISA-seq reads mapping to AAV-Donor. A significant number of these alignments showed AAV-Donor sequences ligated at the 3’ end to another AAV-Donor These reads were a result of vector circularization and tail-to-head concatamerization of AAV-Donor as depicted in **b**. **b** Created with BioRender.com.

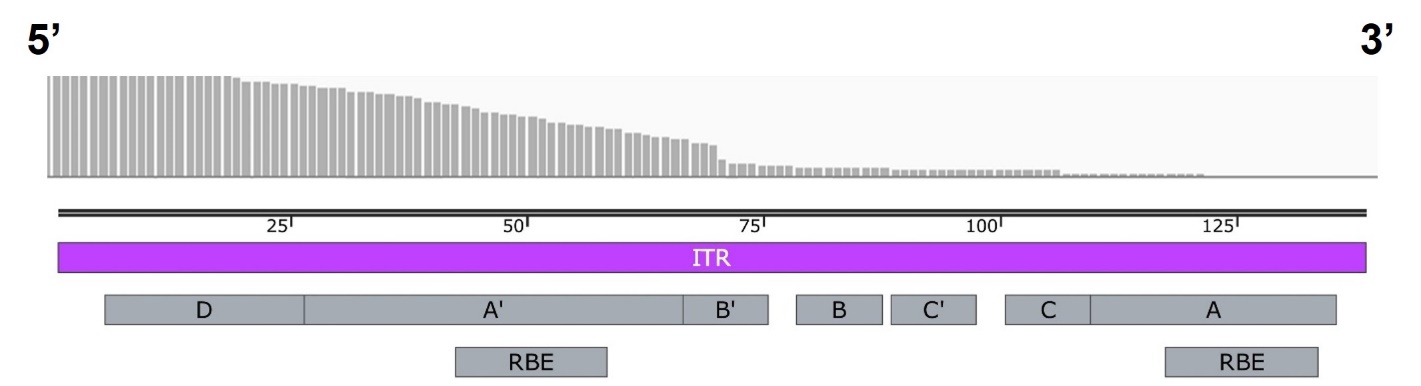

**Supplementary Fig. 20. ITR preferred breakpoint of AAV-Donor integrated in mouse genome.** Enrichment primers used in GISA-seq bind internally to ITR on 3’ end of AAV-Donor, providing full sequence of integrated ITR. We observed a significant drop in coverage between the palindromic B–B′ short arm and A–A′ long arm of the AAV ITR hairpin, indicating a favored breakpoint as previously reported.^3^ RBE: Rep binding element.

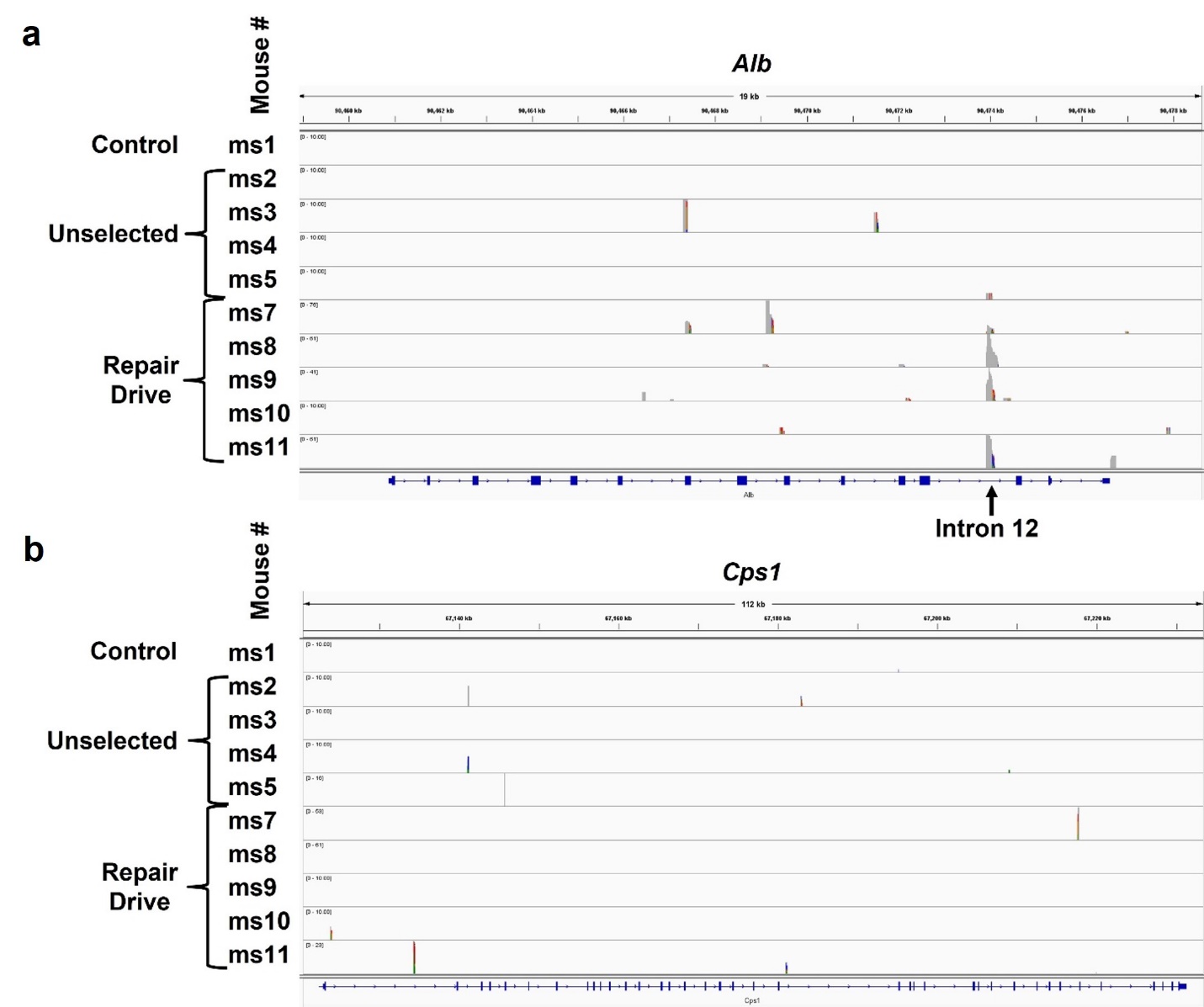

**Supplementary Fig. 21. GISA-seq identified AAV-Donor integration sites in *Alb* and *Cps1*.** IGV visualization of GISA-seq reads mapping in the *Alb* (**a**) and *Cps1* (**b**) locus in Control (ms1), Unselected (ms 2-5) and Repair Drive (ms 7-11) mice. There are not aligned read peaks in Control mouse (ms1) at both the *Alb* and *Cps1* locus. Unselected and Repair Drive mice showed different integration sites in *Alb* and *Cps1*. Interestingly, an integration site in intron 12 of *Alb* was shared between five mice suggesting to be an integration hotspot for AAV-Donor. At the *Alb* locus, integrations occurred at 1-2 peaks in Unselected mice and increased to 2-4 peaks in Repair Drive mice with a higher count at each site, suggesting that they could be originated from clonal expansion of cells harboring those integration events.

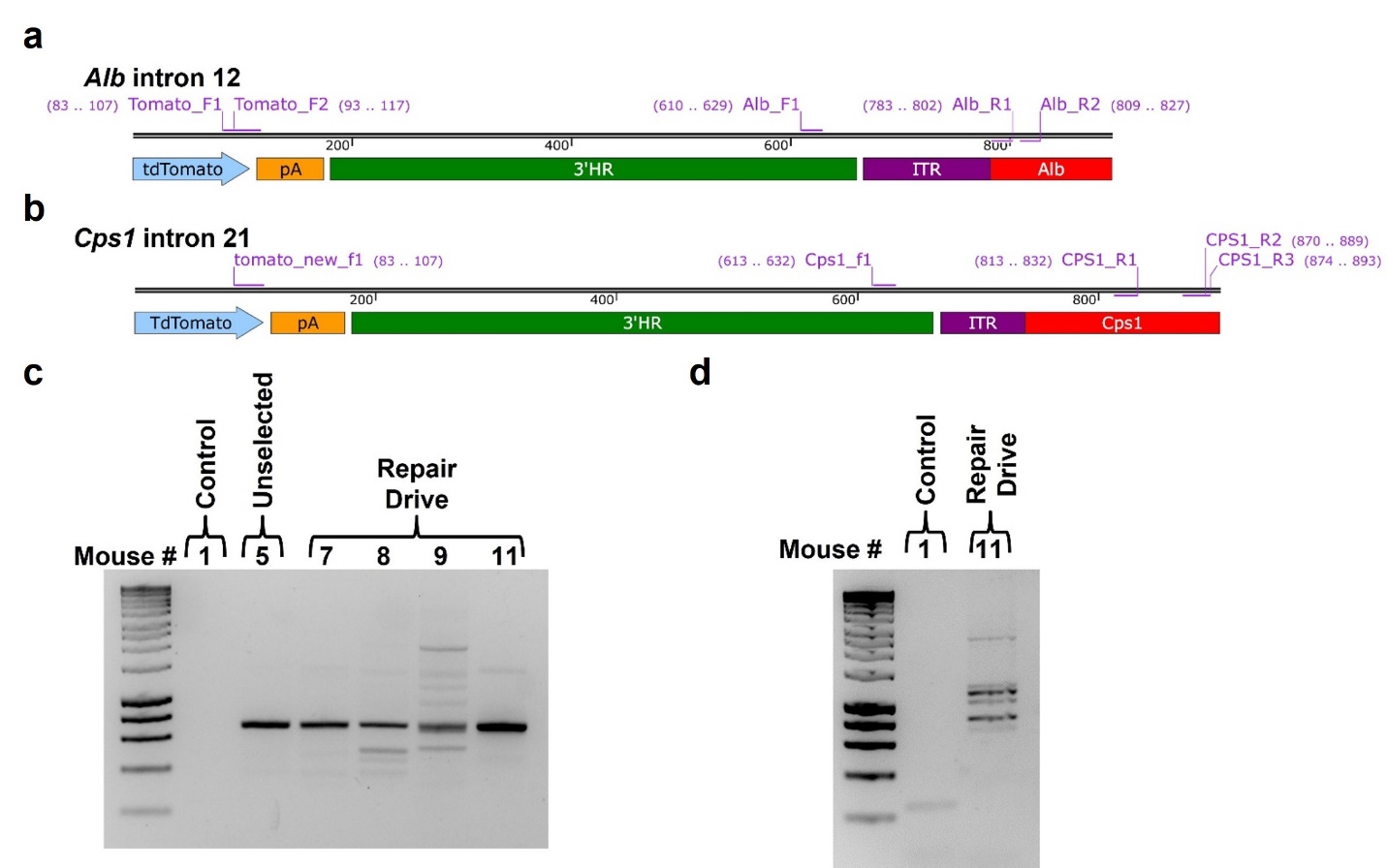

**Supplementary Fig. 22. Confirmation of AAV-Donor integration sites by PCR and Sanger sequencing.** **a**, Primer design to amplify AAV-Donor integrated into intron 12 of *Alb*. **b**, Primer design to amplify AAV-Donor integrated into intron 21 of *Cps1*. **c**, Agarose gel image of PCR products generated using primers amplifying across AAV-Donor integration junction at *Alb.* Amplicon in mouse 11 sample was sent for Sanger sequencing. Expected sequence of TdTomato-3’HR-ITR-*Alb* was confirmed. **d**. Agarose gel image of PCR products generated using primers amplifying across AAV-Donor integration junction at *Cps1*. Amplicon in mouse 11 sample was sent for Sanger sequencing. The expected sequence of TdTomato-3’HR-ITR-*Cps1* was confirmed.

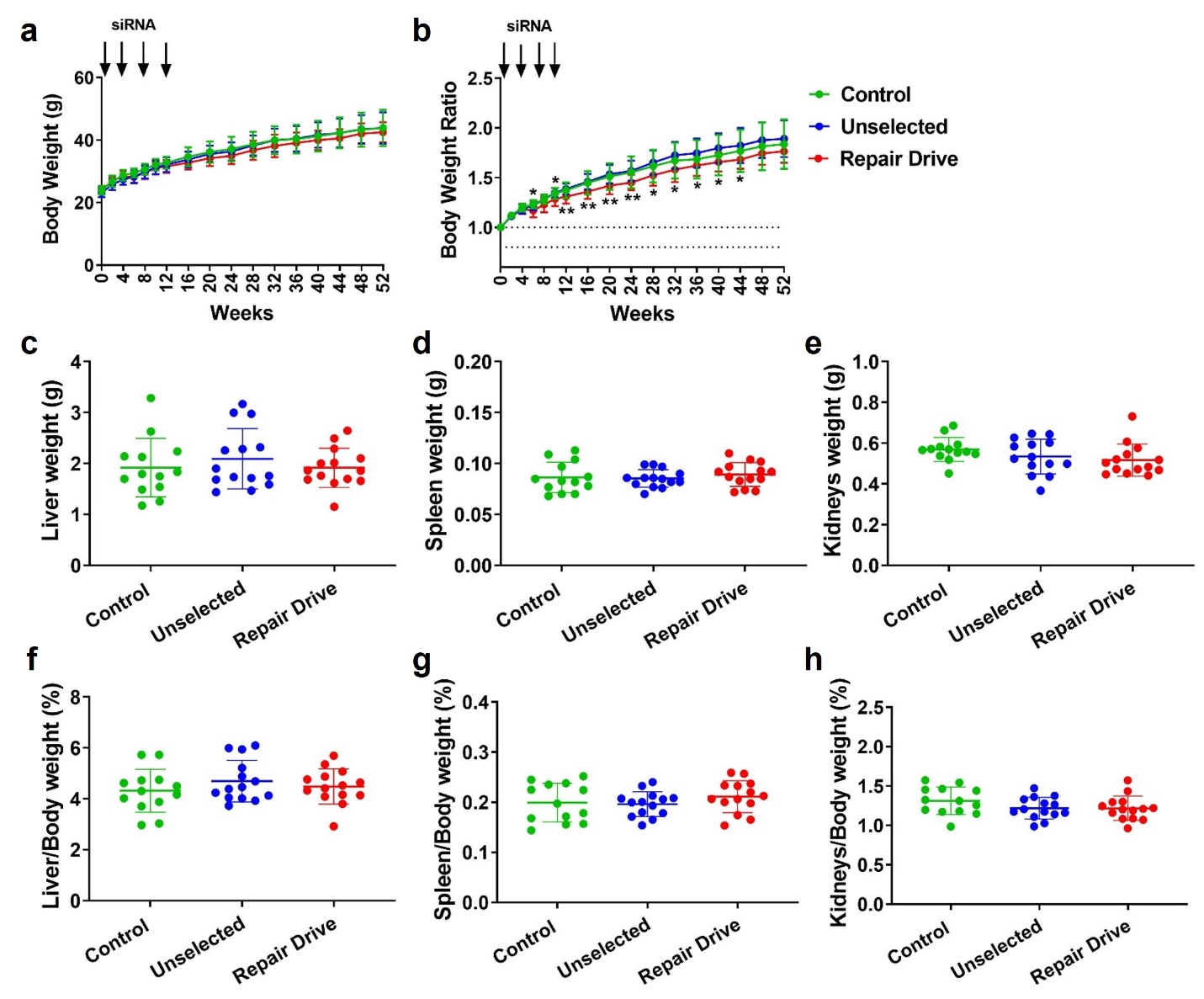

**Supplementary Fig. 23. Body and organ weights at 1 year following Repair Drive.** Body weights (**a**) and body weight ratios normalized to time 0 (**b**) over time. Endpoint weights of liver (**c**), spleen (**d**) and kidneys (**e**). Endpoint organ to body weight ratios for liver (**f**), spleen (**g**) and kidneys (**h**). Data are expressed as mean ± standard deviation (n= 13-14 mice per group), with significance determined by one-way and two-way ANOVA followed by Tukey test, respectively in **c**-**h** and **a**-**b**. In (**b**), * p<0.05 Repair Drive vs. Control at 6 weeks; * p<0.05 Repair Drive vs. Unselected at 10 weeks; ** p<0.01 Repair Drive vs. Unselected at 12, 16, 20 and 24 weeks; * p<0.05 Repair Drive vs. Unselected at 28, 32, 36, 40 and 44 weeks.

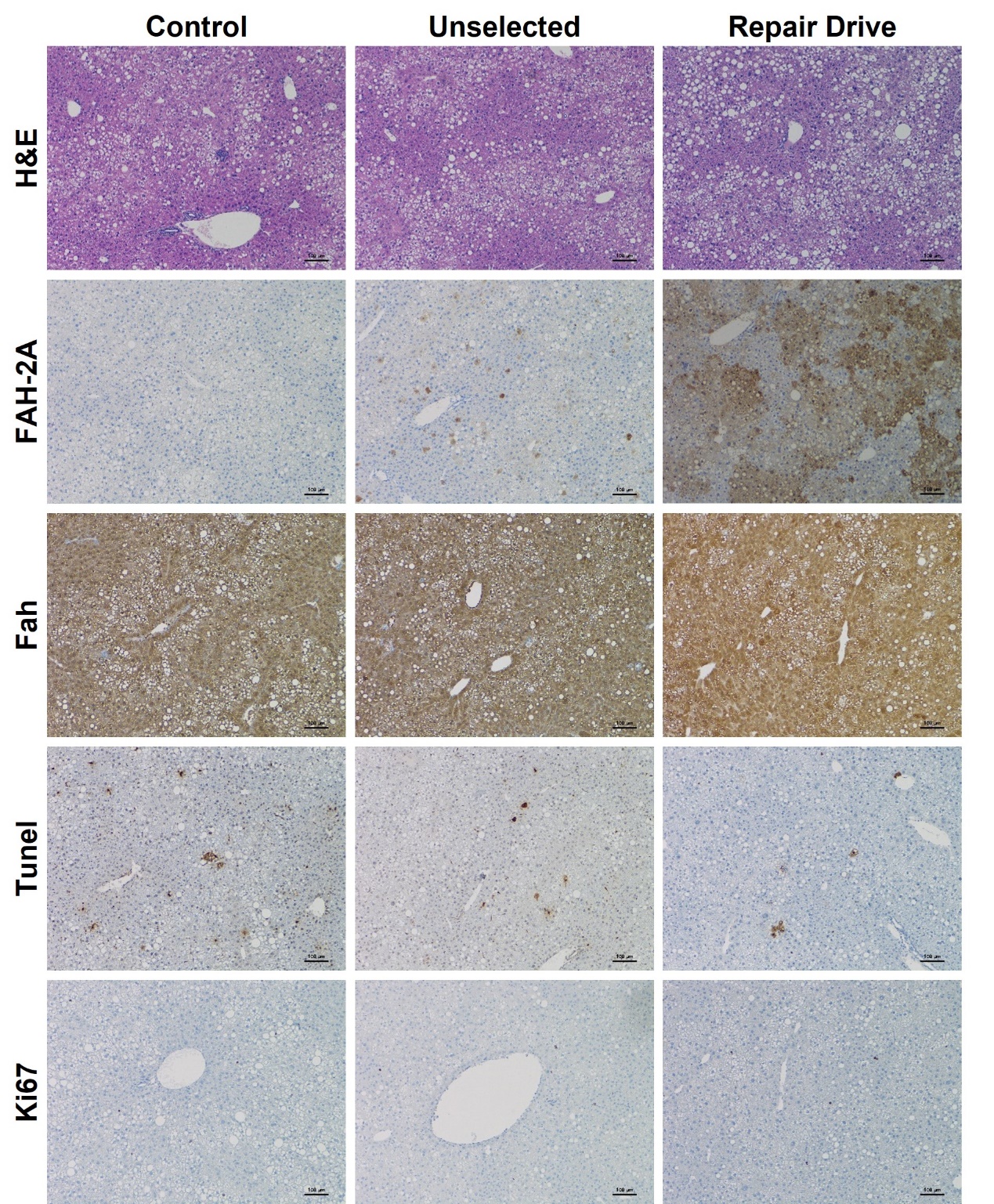

**Supplementary Fig. 24. Representative fatty livers in mice enrolled in the long-term expansion experiment (1 year).** Representative H&E and immunohistochemistry staining of FAH-2A, Fah, Tunel and Ki67 showing liver steatosis in sixty-week-old Control, Unselected and Repair Drive mice. Scale bar is 100 µm.

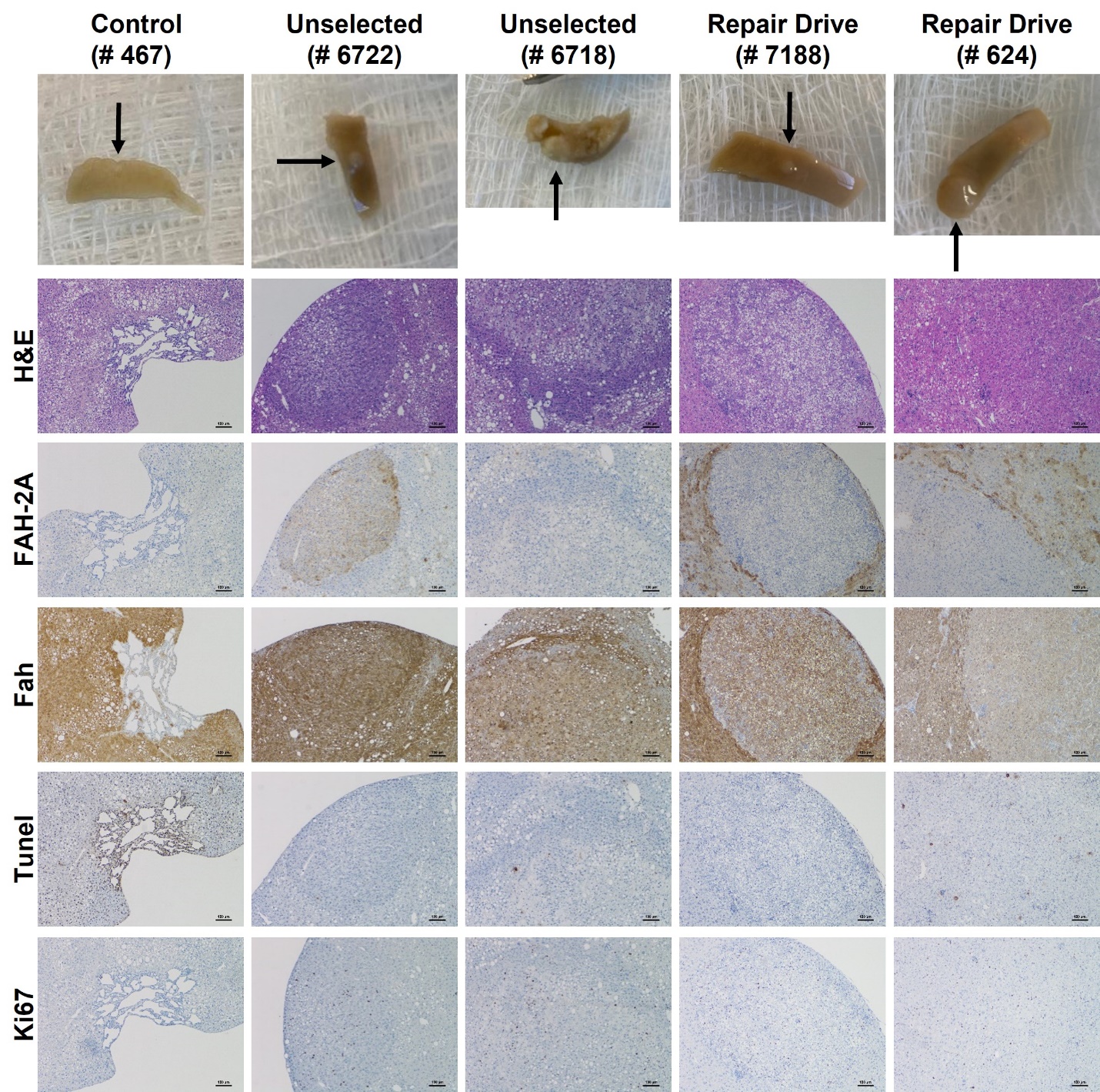

**Supplementary Fig. 25. Liver abnormalities observed in mice enrolled in the long-term expansion experiment.** Photograph, H&E and immunohistochemistry staining of FAH-2A, Fah, Tunel and Ki67 of hepatic abnormalities found in 60-week-old mice (1 year following AAV injection). A hemangioma was found in a Control mouse (# 467), and localized proliferative lesions were found in two Unselected mice (# 6722 and 6718) and two Repair Drive mice (# 7188 and 624). Scale bar is 100 µm.

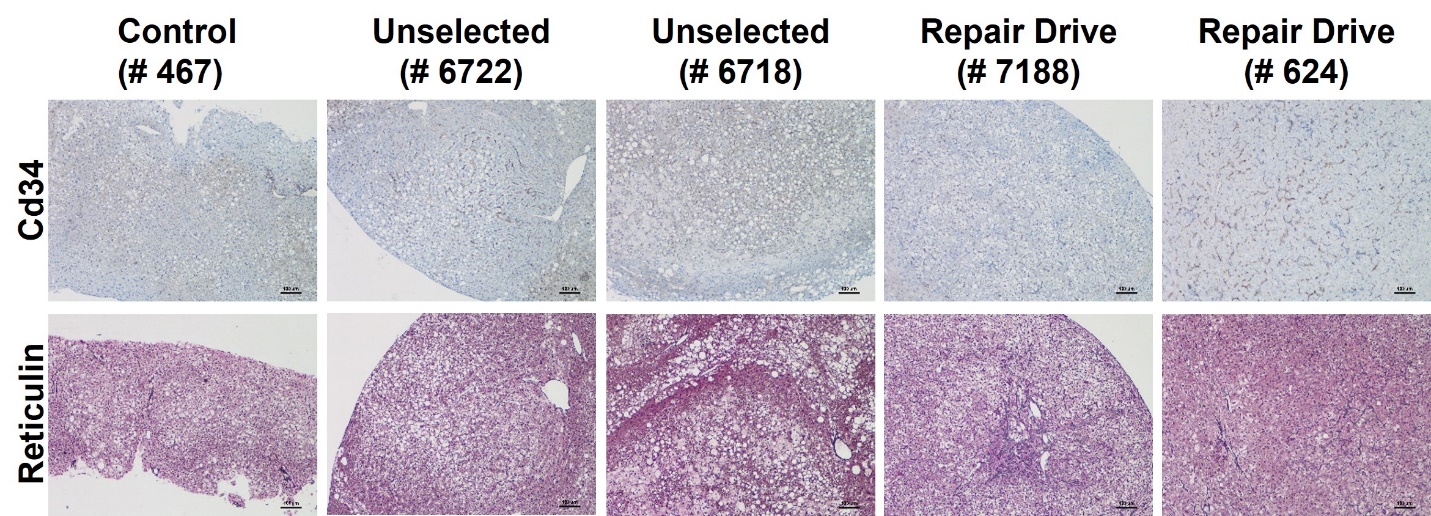

**Supplementary Fig. 26. Characterization of liver abnormalities.** Cd34 and Reticulin staining of hepatic abnormalities found in 60-week-old mice (1 year following AAV injection). Scale bar is 100 µm.
